# Pelagic larval polyclads that practice macrophagous carnivory

**DOI:** 10.1101/2021.02.03.429620

**Authors:** George von Dassow, Cecili Mendes

## Abstract

We report evidence that hatchling polyclads of several genera feed in the plankton on large prey. These ciliated swimmers, despite apparently lacking means to concentrate food or even detect it at a distance, subdue and consume fast-moving active-swimming plankters such as crustacean larvae and copepods, or molluskan veligers. We describe feeding events in captivity using videomicroscopy, and identify several wild-caught predatory pelagic polyclad larvae to genus or species level by DNA barcoding. Remarkably, one of these types is identified unambiguously with a species previously observed as Müller’s larvae, which live as conventional planktotrophs on an inferred diet of small phytoflagellates. Therefore we conclude first that while so-called “direct-developing” polyclad flatworms may hatch with juvenile-like morphology, at least some of these are functionally larvae. Second, that some species of polyclad have at least a triphasic life cycle, with a first larval stage living in the plankton on primary producers followed by a second larval stage living in the plankton by macrophagous carnivory, before presumably settling to the benthos for adult life.

## 1. Introduction

Polyclad flatworms are spiralian lophotrochozoans that dwell in all marine habitats and mostly live by actively hunting and consuming other animals, including both sessile and motile prey. Several polyclads are known to develop via long-lived pelagic larvae that feed in the plankton before undergoing metamorphosis to benthic, crawling worms (reveiwed by Rawlinson, 2014). The most familiar of these is Müller’s larva (Müller, 1850), which develops multiple paired ciliated tentacles and captures small particulate food using a combination of ciliary and muscular action (von Dassow & Ellison, 2020). Other polyclads lay eggs that develop directly into ciliated hatchlings resembling miniature adults (Shinn 1987; Martín-Durán and Eggers, 2012; Rawlinson 2014). These often swim rather than crawling in a manner befitting the adult (Shinn, 1987; Morita et al., 2018), suggesting that they may represent a dispersal stage, yet their way of making a living or transitioning to adult habitat and behavior seems largely unknown.

The small size of many direct-developing polyclad hatchlings raises the question, how and what do they eat? Since some adult polyclads are suctorial feeders, removing small portions of prey near the surface, juveniles of these species conceivably feed similarly as long as prey access poses no challenges. Likewise for species that live by scavenging. For other polyclads that, as adults, specialize on live and motile or large and well-defended prey, it seems likely that small offspring must occupy a distinct niche.

Here we describe macrophagous carnivory by a group of pelagic polyclad “juveniles”. The most prevalent of these small ciliated swimmers capture planktonic crustaceans, mostly larvae, and devour them whole or nearly so by inserting the pharynx beneath the cuticle and ingesting soft tissue from living prey. Other morphotypes capture and devour gastropod or bivalve veligers. These worms lack any special traits that distinguish them from their parents, hence are apparently direct developers. Yet, because they apparently feed and grow in a habitat distinct from their parents, these are functionally larvae. DNA barcoding shows that all of these are acotyleans, many related to the genus *Notocomplana*, which are otherwise categorized as direct developers. Others correspond to unknown acotylean species previously identified as Müller’s larvae.

## 2. Methods

Pelagic flatworms and their prey were collected in regular dockside plankton tows in the Charleston Marina using a 53-micron mesh net with a jar cod end during January-May 2020. The majority of samples were taken shortly before the high slack. Concentrated plankton was diluted with an approximately equal volume of filtered natural seawater, and samples for sorting were taken over the course of several hours from the surface on the window side of the jar. Specimens were isolated by hand selection into bowls of filtered natural seawater and kept in sea-tables at ambient seawater temperature of 12-15 °C. Prey was isolated similarly from natural plankton, or was obtained by shining a desk lamp onto one end of a seatable housing barnacle-covered rocks (*Balanus glandula*). Adult flatworms (tentatively identified as *Notocomplana litoricola*) were collected from Fossil Point (GPS 43.3575, -124.3118) or Lighthouse Beach (GPS 43.3400, -124.3749), Coos Bay, Oregon from underneath high-intertidal rocks, then housed in clear plastic screw-cap pint jars (Talenti Gelato) with daily water changes and small barnacles (*Balanus glandula*) for food. Egg plates were deposited promptly and continuously after collection.

All microscopic observation and video recording was conducted at ambient room temperature of 16-18 °C. Recordings of feeding were performed either in Syracuse dishes on a Leica Z6 Apo macroscope or on slide-and-coverslip cuvettes on an Olympus BX51 equipped for DIC. Video and time-lapse recording was achieved with a Point Grey Grasshopper 3 camera operated by StreamPix 7, which enables recording frames at variable rate into a circular buffer. Figure assembly was performed using ImageJ and Adobe CS6. No convolution filters were applied to any image except as noted explicitly in the legend, and all panels preserve original handedness.

For DNA barcoding, planktonic specimens were isolated without food for one day or more, then washed by allowing them to swim briefly in fresh filtered seawater, before placing individually in microfuge tubes with <10 µl seawater and frozen at -80 °C. Adult worms were rinsed, then placed on a clean plastic petri dish in a slick of filtered seawater. A posterior fragment was severed with a clean razor blade, and transferred to a microfuge tube in a minimal drop of water, then frozen at -80 °C. Genomic DNA from planktonic specimens was extracted with Chelex matrix (InstaGene; BioRad) and from adult tissue with DNEasy Blood and Tissue kit (Qiagen), using manufacturer instructions. The barcode regions of mitochondrial cytochrome c oxidase subunit 1 (COI) and 16S ribosomal RNA (henceforth, 16S rRNA) were amplified using the primer pairs LCO1490/HCO2198 (Folmer *et al*. 1994), ar-L/br-H (Palumbi *et al*. 1991), and universal variants jgLCO1490/jgHCO2198 (Geller *et al*. 2013), and using the crustacean specific primer pair COI6Lb/COIAH2m (Schubart & Huber, 2006; Mantelatto *et al*. 2016) to amplify possible gut contents. Polymerase chain reactions (PCR) were carried out using GoTaq Green Master Mix (Promega) in the following conditions: initial denaturation at 94 °C for 1 min; 35 cycles of denaturation at 94 °C for 30 s; annealing at 43°-45 °C (COI) or 47-50 °C (16S rRNA) for 1 min, extension at 72 °C for 1 min; and final extension at 72°C for 5 min. PCR products were purified with Wizard SV Gel and PCR Cleanup kit (Promega). Purified products were sequenced at Sequetech DNA Sequence Service (Mountain View, CA-USA). Consensus sequences were obtained using GeneStudio ™ Pro (GeneStudio, Inc.), and COI sequences were checked for stop codons. Resulting sequences were deposited in Genbank (Table 1). Sequences were aligned in the online version of Mafft software (Katoh et al. 2019), and analyzed using Automatic Barcoding Gap Discovery (ABGD) online software (Puillandre et al. 2012) with default values for all parameters. The resulting sequences were also used as input for BLASTn through the NCBI website and the five closest matches to each were incorporated to the dataset. All sequences were aligned in Mafft and used as input for phylogenetic inference in RAxML v.8.2.12 (Stamatakis 2014), as available in Cipres (Miller et al. 2010), under GTRGAMMA model with 1,000 bootstraps, with *Imogine* cf. *aomori* as root for COI and *Imogine fafai* for 16S rRNA. The resulting trees were visualized in FigTree v.1.4.3 (Rambaut, 2014).

**Table 1.** Pelagic flatworms with accession numbers, OTUs, and closest match in GenBank. Corresponding trees in Fig. 15.

| Specimen identifier | Operational Taxonomic Unit (OTU) | Closest GenBank Match (percentage of similarity) | GenBank Accession 16S rRNA | GenBank Accession COI |
| --- | --- | --- | --- | --- |
| COIMB2020_PFW03 | Notocomplana sp. 2 | MF398458 - Notocomplana palta (94.57%) | MW399190 | — |
| COIMB2021_PFW04 | Notocomplana sp. 2 | MF398458 - Notocomplana palta (94.83%) | MW399191 | MW384675 |
| COIMB2022_PFW05 | Kaburakia excelsa | KT363735 - Holoplana (89.78%) | MW399207 | — |
| COIMB2023_PFW06 | Notocomplana sp. 1 | MF398458 - Notocomplana palta (93.88%) | MW399199 | — |
| COIMB2024_PFW07 | Notocomplana sp. 2 | MF398458 - Notocomplana palta (94.65%) | MW399193 | MW384676 |
| COIMB2025_PFW08 | Notocomplana sp. 2 | MF398458 - Notocomplana palta (94.65%) | MW399195 | — |
| COIMB2027_PFW10 | Notocomplana sp. 2 | MF398458 - Notocomplana palta (94.48%) | MW399196 | — |
| COIMB2028_PFW11 | Notocomplana sp. 1 | MF398458 - Notocomplana palta (93.29%) | MW399201 | MW384681 |
| COIMB2029_PFW12 | Notocomplana sp. 2 | MF398458 - Notocomplana palta (94.22%) | MW399194 | MW384680 |
| COIMB2030_PFW13 | Notocomplana sp. 2 | MF398458 - Notocomplana palta (94.32%) | MW399198 | MW384677 |
| COIMB2031_PFW14 | Notocomplana sp. 2 | MF398458 - Notocomplana palta (94.96%) | — | MW384678 |
| COIMB2032_PFW15 | Notocomplana sp. 2 | MF398458 - Notocomplana palta (94.87%) | MW399192 | — |
| COIMB2033_PFW16 | Notocomplana sp. 1 | MF398458 - Notocomplana palta (93.82%) | MW399200 | — |
| COIMB2034_PFW17 | Notocomplana sp. 2 | MF398458 - Notocomplana palta (94.9%) | MW399197 | MW384679 |
| COIMB2035_PFW18 | Notocomplana litoricola | LC176061 - Notocomplana setentrionalis (94.04%) | MW399202 | — |
| COIMB2037_PFW20 | Unknown Acotyela | MN017755 - Acotyela sp. (100%) | MW399210 | — |
| COIMB2038_PFW21 | Unknown Acotyela | MN017755 - Acotyela sp. (100%) | MW399211 | — |
| COIMB2039_PFW22 | Unknown Acotyela | MN017755 - Acotyela sp. (100%) | MW399215 | — |
| COIMB2045-PFW23 | Kaburakia excelsa | KT363735 - Hoploplana elisabelloi (89.88%) | MW399208 | MW384684 |
| COIMB2050-PFW24 | Notocomplana sp.3 | LC176050 - Notocomplana koreana (93.06%) | MW399206 | — |
| COIMB2052-PFW26 | Kaburakia excelsa | KT363735 - Hoploplana elisabelloi (90.37%) | MW399209 | MW384685 |
| COIMB2054-PFW28 | Unknown Acotyela | MN017755 - Acotyela sp. (100%) | MW399212 | MW384687 |
| COIMB2055-PFW29 | Unknown Acotyela | MN017755 - Acotyela sp. (100%) | MW399213 | MW384686 |
| COIMB2056-PFW30 | Unknown Acotyela | MN017755 - Acotyela sp. (99.78%) | MW399214 | MW384688 |
| COIMB2057-PFW31 | Unknown Acotyela | MN017755 - Acotyela sp. (99.78%) | MW399216 | MW384689 |
| COIMB2058-Noto#2 | Notocomplana litoricola | LC176061 - Notocomplana setentrionalis (94.48%) | MW399203 | MW384682 |
| COIMB2060-Noto#1 | Notocomplana litoricola | LC176061 - Notocomplana setentrionalis (94.48%) | MW399204 | MW384683 |
| COIMB2061-Noto#3 | Notocomplana litoricola | LC176061 - Notocomplana setentrionalis (94.27%) | MW399205 | — |
| COIMB2118_PFW41 | Notocomplana sp. 1 | MF398458 - Notocomplana palta (93.64%) | MW553724 | MW556700 |
| COIMB2119_PFW42 | Notocomplana sp. 2 | MH242936 - Polycladida sp. BMBM-0249 (99.24%) | — | MW556701 |

## 3. Results

### 3.1 Small predatory flatworms found in the plankton eat crustacean larvae

Apparently-pelagic small flatworms were recovered from nearly all dockside plankton samples. By far the most common was a kind with multiple pairs of central eyespots, posterior mouth, and a characteristic set of marginal cirri. The set included regularly spaced anterior marginal cirri, a widely-spaced lateral pair, and a prominent posterior pair (Fig. 1). Individuals in plankton samples (before feeding in captivity) were as small as ∼150 µm and as large as 1 mm, and varied in pigmentation and the presence and size of spherules in digestive diverticula.Figures 2 and 3 depict two additional distinctive morphotypes that were less frequent but still encountered repeatedly: one with bright red pigment (everywhere in small specimens, in anterior and posterior patches in larger ones) and a closely-spaced marginal pair of cirri, and another with very prominent, large, and numerous spherules in the gut. For all three of these types, small individuals were oval, tending toward keyhole-shaped (larger anterior), whereas larger ones tended to hold themselves in a kite shape. The smallest had two pairs of eyespots, and the largest when caught had more, up to eight pairs, some cerebral and some nuchal. When kept in bowls, both large and small animals underwent active ciliary swimming in addition to surface gliding, and were also frequently seen at rest on the bowl surface or on the water surface. When gliding near a surface, they always put mouth toward surface and moved forward. They were not sticky at this stage, exhibited pronounced metachronal waves of ciliary activity across most of their surface, and did not appear to leave significant mucus trails.

**Figure 1.**
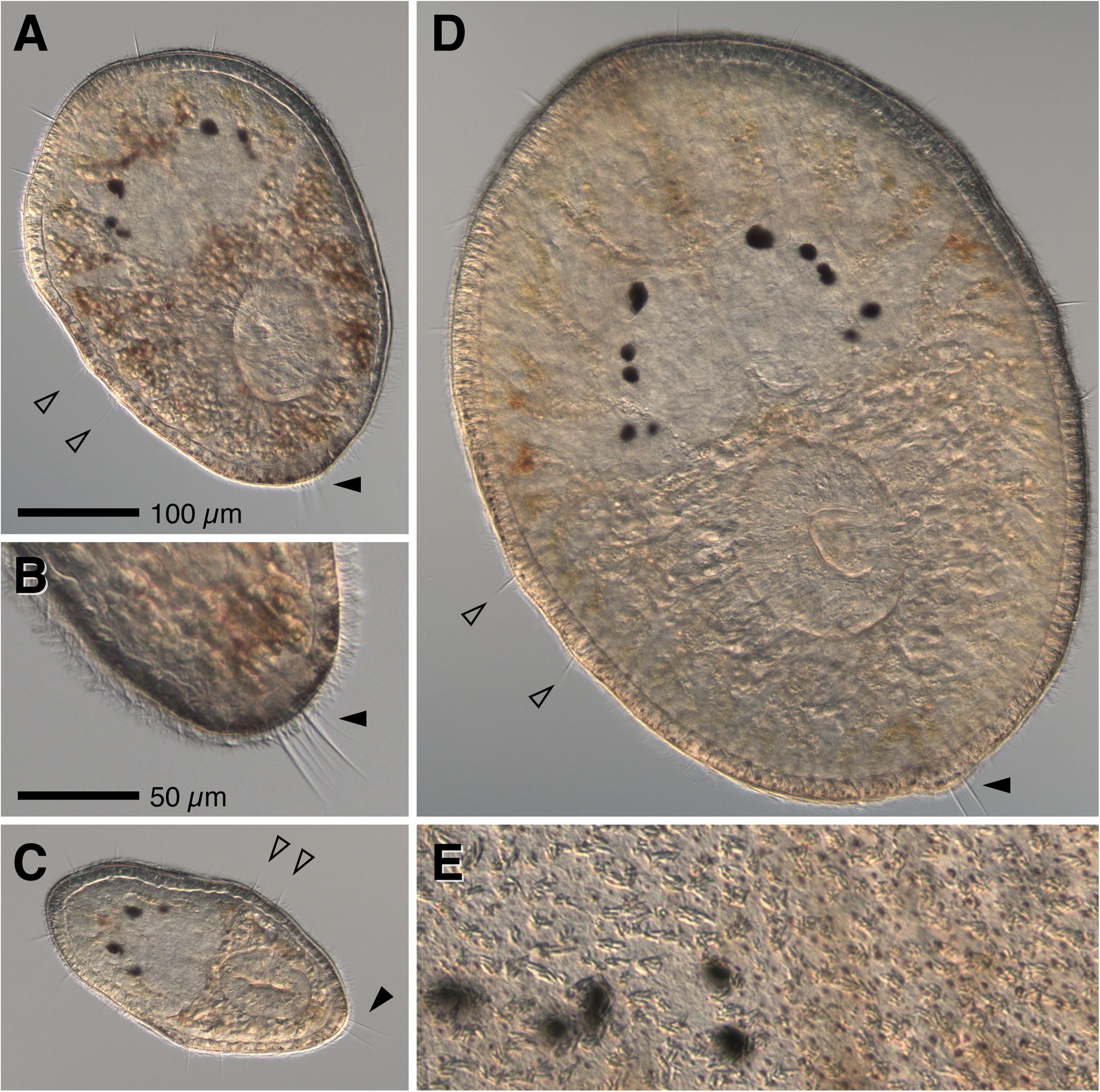
The most abundant variety of nauplius-eating pelagic flatworm. (A) Typical specimen shortly after collection, ∼300 µm and only faintly colored, with four pairs of eyespots. Note the regularly-spaced anterior marginal cirri (six are visible here), lateral pair of well-separated discrete cirri (hollow arrowheads), and posterior cirrus pair (solid arrowhead; out of focus). (B) 2x blow-up showing the posterior end of the same individual, showing that the posterior cirrus pair is surrounded by numerous long non-beating cilia. (C) Small individual of the same type, with two pairs of eyes and identical cirrus arrangement. (D) Large individual of the same type, five eyespot pairs, same cirrus arrangement. The posterior two pairs of eyespots are just under the epidermis; these become nuchal eyespots, although tentacles do not protrude until much later, and then only barely above the surrounding epidermis. (E) Superficial focus on a different specimen, similar in size to D, showing clustered rhabdites in the epidermis and higher density of small freckles posterior to the nuchal eyespots.

**Figure 2.**
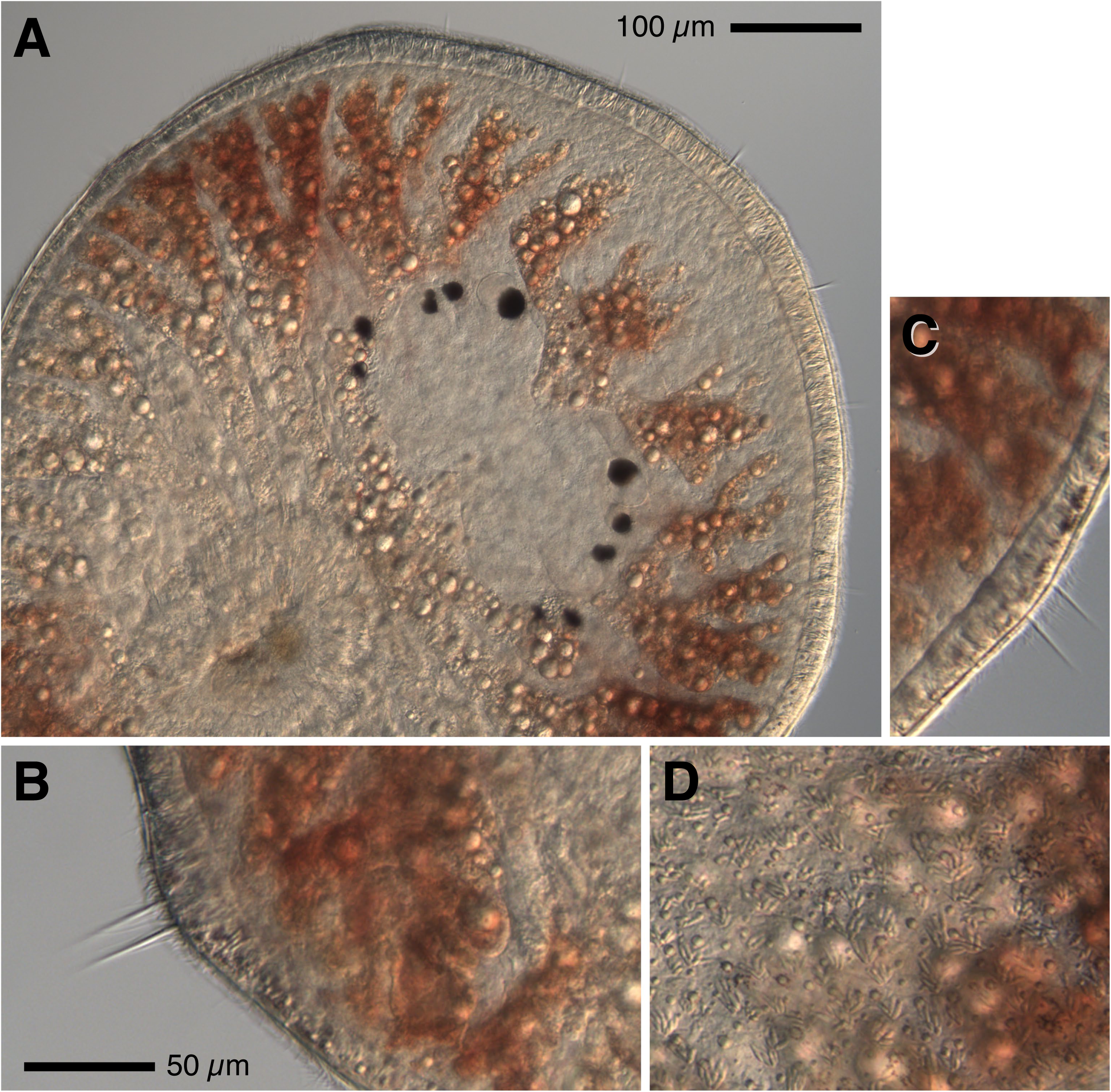
The red-cheeked nauplius-eater. (A) A large specimen with deep red pigmentation around anterior and posterior, but not midsection, digestive diverticula. (B-D) 2x blow-ups of posterior, lateral, and superficial patches on the same individual. These have nearly the same cirrus arrangement as the type depicted in Figure 1 – six anterior, two lateral, paired posterior with accessory cilia – but as shown in C, the lateral cirri are closely paired.

**Figure 3.**
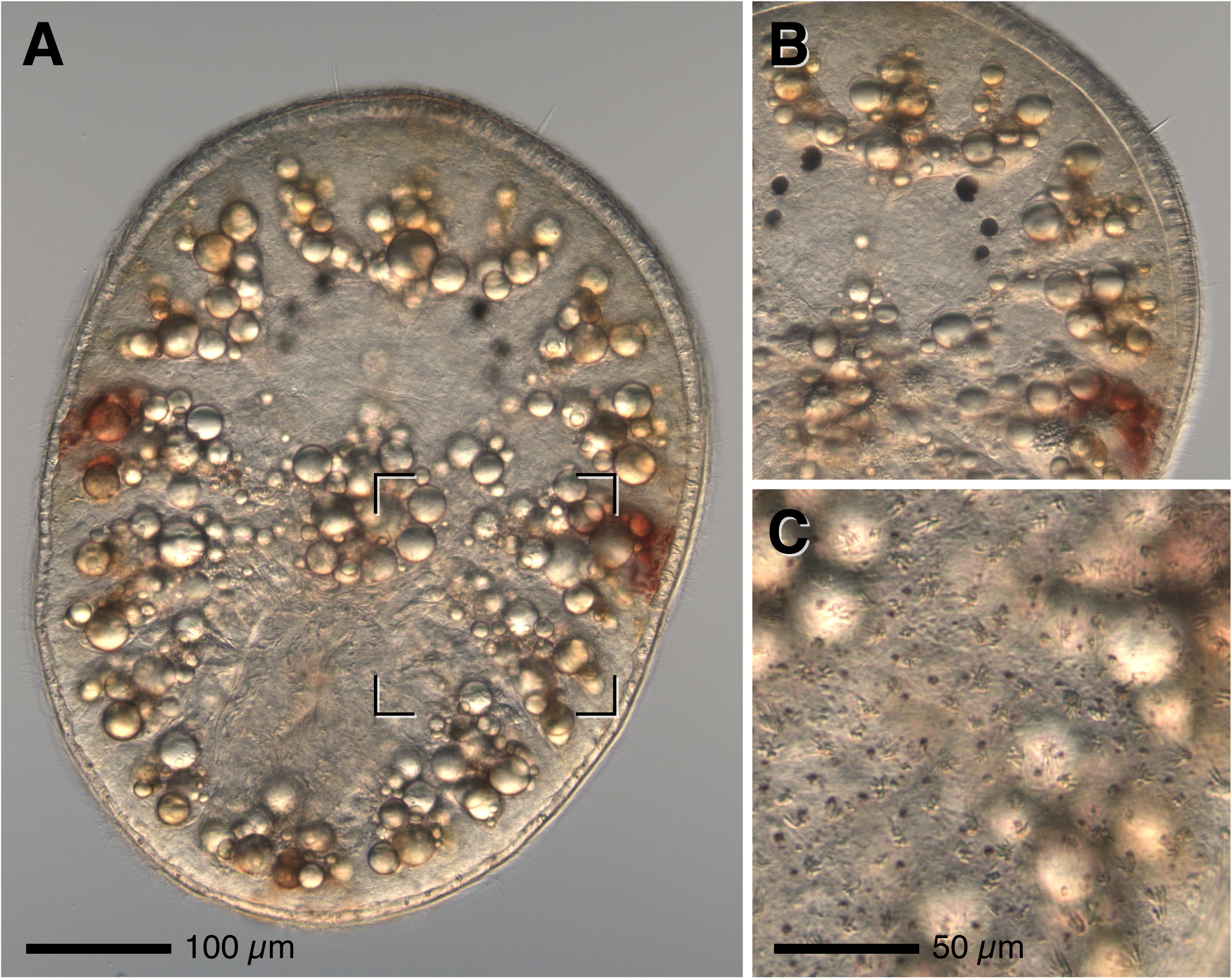
A third nauplius-eating morphotype. (A) Focal plane dominated by large refractile spherules in the digestive diverticula. These decrease with starvation but do not appear to represent direct food remains, which are cloudy and shapeless. (B) Focal plane in the same individual, with eyespots in focus. (C) 2x blow-up of a superficial view in the same individual, at the position shown by brackets in (A), illustrating freckles posterior to the brain. Most individuals of this type also have small red patches laterally or posteriorly (as this one does) and faint yellow pigmentation elsewhere.

These flatworms were initially collected out of curiosity, and housed with other plankters of interest in common bowls containing several dozen animals of various types, especially hoplo- and palaeonemerteans and their likely prey, which included molluscan veligers, polychaete setigers and nectochaetes, and crustacean nauplii. While surveying such bowls by stereomicroscope, the pelagic flatworms were observed attaching to, occasionally riding or swimming with, and then devouring barnacle nauplii. Thereafter, pelagic flatworms of this general type were isolated and offered various potential crustacean prey including barnacle or copepod nauplii, copepodites and adult copepods, and decapod zoeas. Although barnacle nauplii were far more often consumed, many instances were observed in which these flatworms consumed copepods or their nauplii, and even decapod zoeas. Observation suggests that these less-frequent prey may be more successful at batting away the flatworm predator through appendage motions or sheer rate of movement.

### 3.2 Prey capture and consumption

Hundreds of feeding events were observed, and several recorded either using video or series of still photographs (Figs. 4, 5, and 6; Supplemental videos 1–4). Feeding began with direct, apparently chance encounters as the paths of slowly-circling worms intersected either swimming nauplii or, more commonly in microscopic observations, nauplii trapped by adhesion to glass. There was no sign that the flatworms sense prey at a distance of more than a body length and direct their movements toward it. However, when approaching within approximately a body length of a moving nauplius, flatworms were often seen to turn sharply toward it. In several instances, one or more worms joined a kill, swimming steadily toward another with either a large prey item or with multiple trapped prey. When feeding on barnacle nauplii, flatworms exhibited no detectable scale preference: large flatworms ate small nauplii, and small ones attacked and ate late-stage nauplii vastly exceeding their own size.

**Figure 4.**
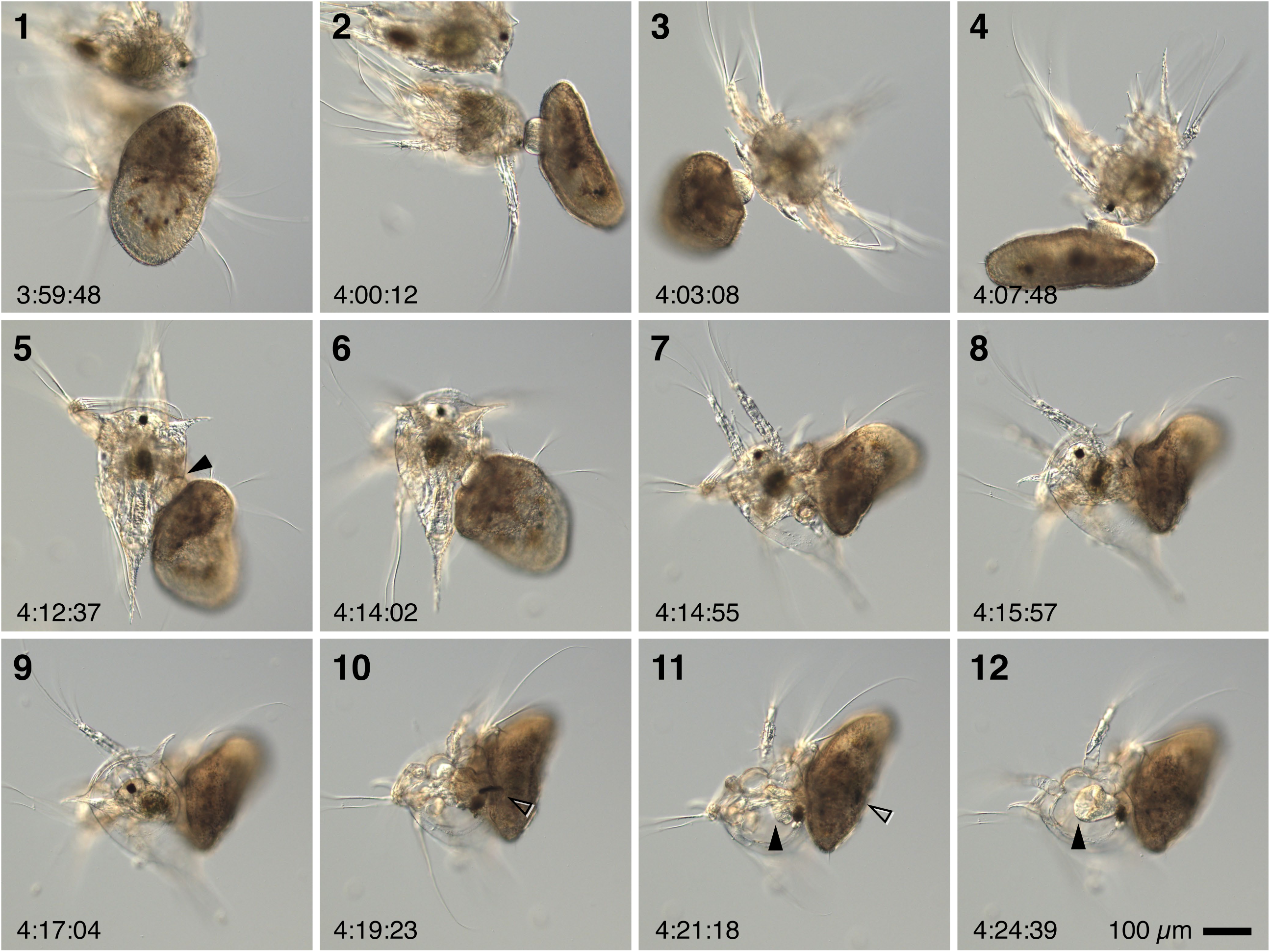
Flatworm captures and eats a barnacle nauplius. Images are taken from a single feeding event recorded in an unconfined chamber as a series of still images (playable sequence in Supplemental Video 1). In frames 1–4, the flatworm attaches by its everted pharynx to the anterior-dorsal portion of the nauplius’ carapace, and swims with its catch. The nauplius is actively attempting to swim too at this stage. In the second row the pair has settled to the bottom of the dish. Frame 5 shows the flatworm’s pharynx (arrowhead) intruded into the nauplius’ armpit. Comparing this zone in frame 6, a clear space has developed, indicating that the worm has begun to swallow prey tissue. In frame 7 the worm has nearly swallowed the entire posterior end, and by frame 8 is beginning to pull limbs from the cuticle. Note that in frames 5–8 the nauplius continues to move its limbs, stroking the first antenna vigorously even while the flatworm engulfs its entire posterior end. In the third row, the flatworm swallows the head and most of the appendages; the naupliar eye (hollow arrowhead) often persists for some time in the worm’s digestive system and provides a visual indication even at low magnification of recent meals. In frames 11 and 12 it is particularly apparent how the flatworm flares out its pharynx (arrowhead) to actively explore the inside of the prey carapace, retrieving many of the stray gobbets that tear off of the main corpus as it is swallowed (note the distal segments of one first antenna remaining, frame 12).

**Figure 5.**
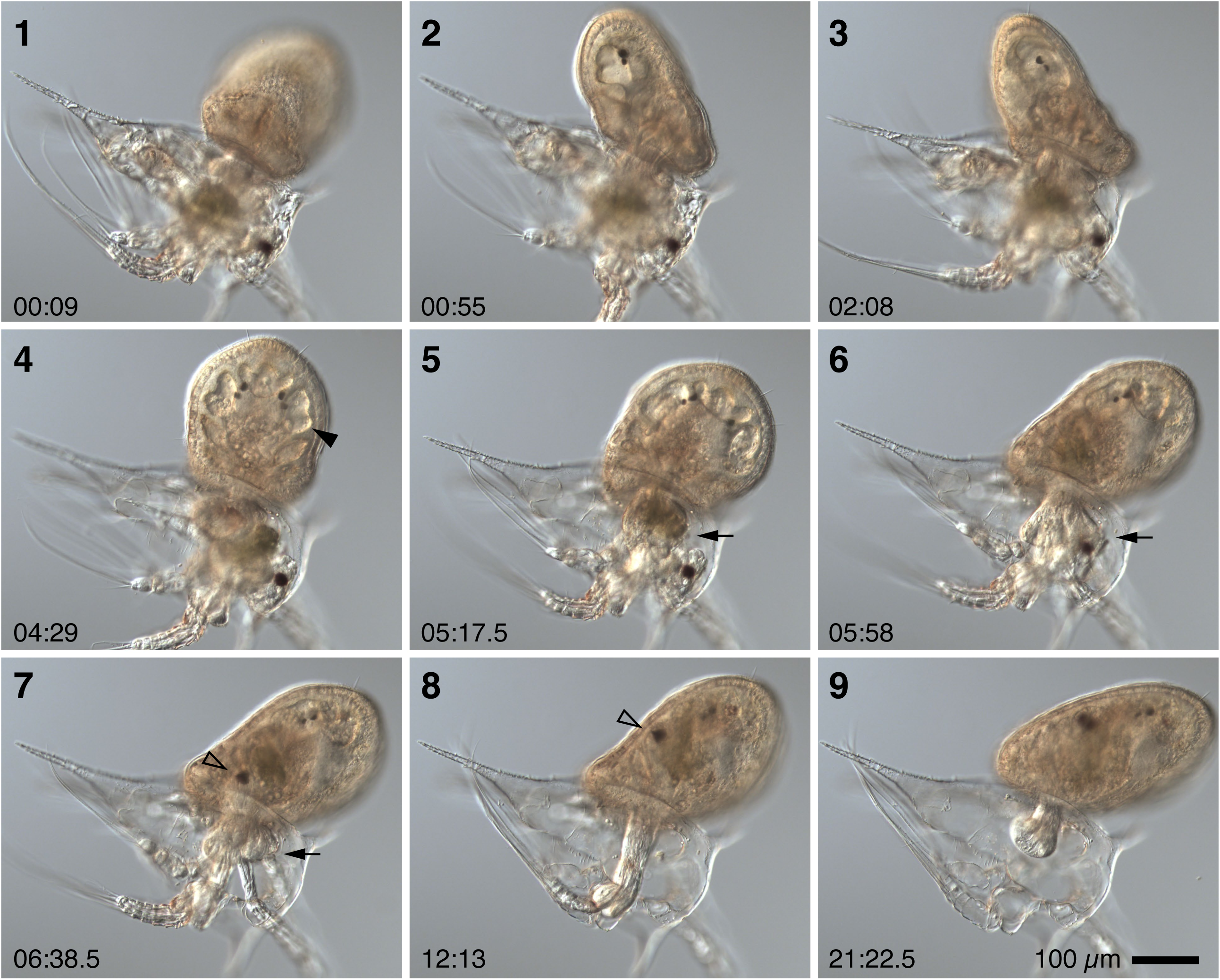
Flatworm captures and eats a barnacle nauplius, time-lapse sequence. Corresponds to Supplemental Video 2. The flatworm had already attached to its victim before they were transferred gently to a slide and coverslipped for recording. The sequence proceeds similarly to Fig. 4 but was recorded continuously. Times in minutes:seconds from the start of recording. The nauplius’ limbs move actively during the first minutes, reducing to twitching as the body is swallowed. In this sequence it is particularly clear that the flatworm inflates its digestive diverticula with fluid (arrowhead, frame 4); this is not always observed, but might either represent preparation for swallowing a large meal or the accidental initiation of pumping prior to actual engagement with prey tissue. Initially the pharynx intrudes just behind the naupliar armpit, but once a significant amount of tissue has been ingested the flatworm extends the pharynx around the remaining corpus (arrow in frames 5–7). The naupliar eye is indicated by a hollow arrowhead in frames 7 and 8 as it is swallowed. This flatworm was remarkably thorough, leaving behind a nearly empty husk. The corresponding video sequence emphasizes the exploratory and prehensile use of the pharynx, which can be seen extending all the way into an appendage in frame 8.

**Figure 6.**
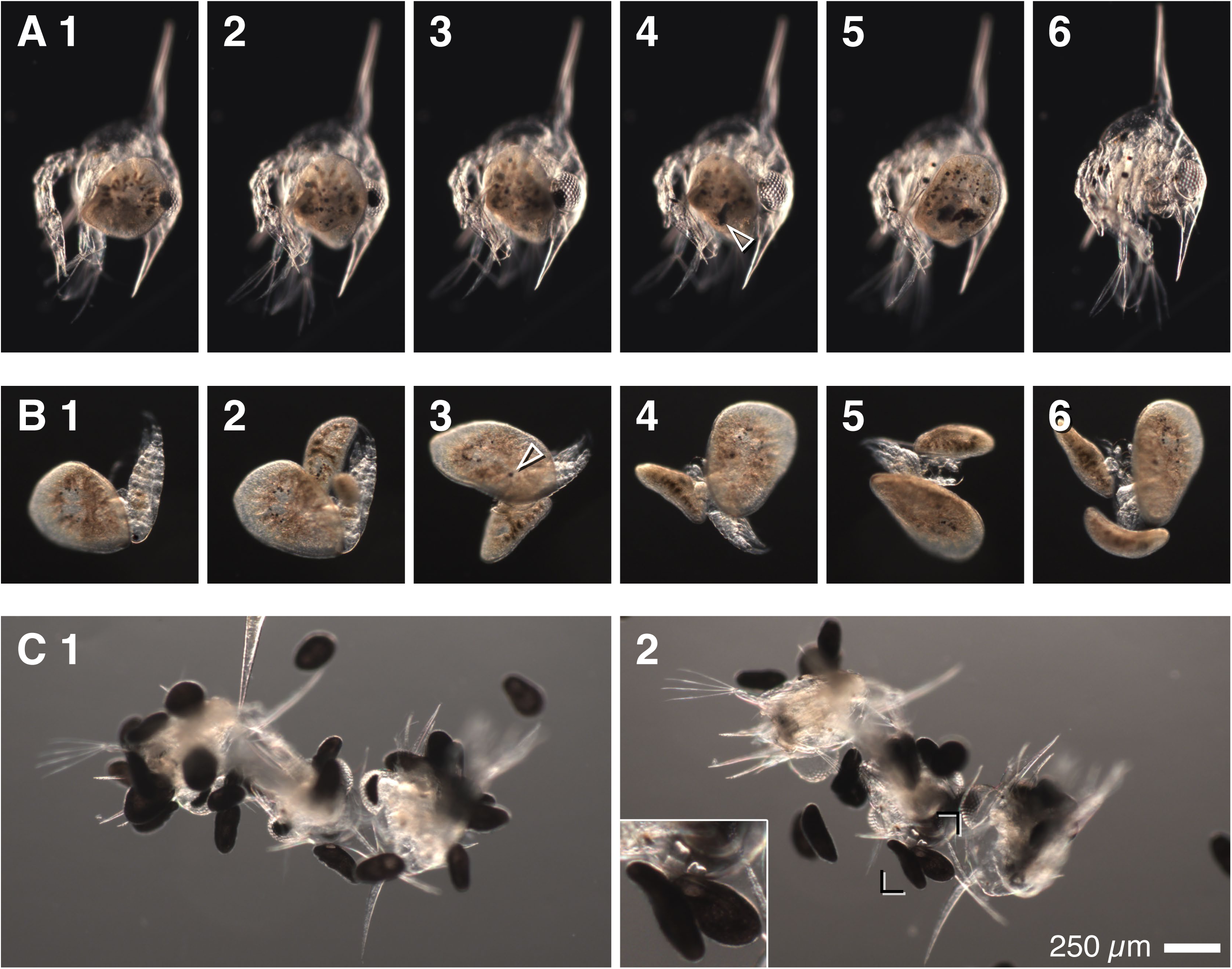
Flatworms prey upon various planktonic crustaceans and feed in groups. (A) A small individual somehow subdues a decapod zoea and eats its head; corresponds to Supplemental Video 3. Prey engagement was not observed in this case. The zoea was initially motile, and attempted to scrape the worm away using its tail, but failed to do so because the worm lodged itself in front of the lateral spine. The worm apparently penetrated the cuticle beneath the carapace, and removed brain and eyes. Arrowhead in frame 4 points out the ingested right eye (note the empty lens). The left followed shortly, whereafter the worm was apparently sated and left the rest of the body twitching feebly. (B) A largish worm devours a small calanoid; corresponds to Supplemental Video 4. This prey was likely captured while attempting to molt. The kill was joined successively by smaller worms. Arrowhead in frame 3 indicates the ingested eye of the copepod. (C) A swarm of several dozen hatchling *Notocomplana litoricola* devours three decapod zoeas; corresponds to Supplemental Video 5. These three prey were initially motile, but were overwhelmed by numerous clinging worms, became entangled together, and were almost completely devoured (note the empty eyesockets in frame 2) over the course of an hour despite the minuscule size of the worms. Inset is a 2x blowup highlighting two individuals side by side with pharynges intruded through a crack in the carapace. Scale in (C) applies to all.

The following description generalizes in present tense from numerous observations. Not all encounters lead to feeding, but in those that do, the first phase – prey engagement – consists of a circling motion around the prey, lasting several seconds to a minute. The worm then everts its pharynx, eventually attaching to the nauplius’ cuticle, often at first on the anterior dorsal carapace (second phase – attachment; Fig. 4, frames 1–4). It seems likely that at this point the worm generally tries to swim off with its prey; in clean bowls or on slide preps, this was often impossible because the prey had become stuck to one or another glass surface. Once the worm and its prey were attached to each other in this manner, it was sometimes possible to transfer predator and prey together by pipette to a slide and coverslip prep without dissociating them.

Initial attachment to the carapace does not in and of itself appear to subdue the prey, but there was often some sign of an effect: naupliar tissue near the attachment site seemed to shrink away from the cuticle. Occasionally, a fortunate view showed that a small hole developed where the worm had attached, suggesting that the worms can penetrate the cuticle. However, in no case was the worm observed to devour its prey through the naupliar carapace. Rather, after the attachment phase, which might last a few minutes, in the third phase – penetration – the worm detaches and moves, without maintaining apparent direct contact, to position its mouth at one of several sites that seem to afford easier access. The most common successful point of entry is near the “armpit” either between the second antenna and mandible or behind the mandible (Fig. 4, frames 5,6). It appears that here the worm can break into a suture between head shield and ventral cuticle, or between body cuticle and appendage.

Once the flatworm penetrates the prey cuticle, it intrudes its pharynx, flared into a trumpet shape, and begins to pump rhythmically (Fig. 4, frames 9–12 and Fig. 5, frames 5–7). During this consummate phase, the worm sucks living, intact prey tissue through the pharynx and deep into digestive diverticula where it very rapidly loses coherence (Supplemental Video 1,2). The prey is still motile at the beginning, and often well beyond, making video recording of non-attached cases nearly impossible; limbs seem to cease thrashing around the time the brain is ingested. Hence it does not appear that the flatworm intoxicates its prey. There was also no definitive sign that the worm injects a digestive aid, as whole tissue masses, including muscle fibers and setal linings, passed the pharynx without dissociation. Even so the prey tissue often tears, leaving limb flesh or body masses behind. These the worm eventually retrieves – unless it detaches upon apparent satiation – during an extended exploratory period, protruding the flared-out pharynx deeply into the emptying cuticle and sucking up as much as it can latch onto (Fig. 5, frames 8,9; note time stamps).

Flatworms often succeed in almost completely emptying the prey, even when it initially exceeds their own linear dimension. Large individuals will proceed immediately to new prey items if available, sometimes consuming several nauplii in the course of an hour or so. Small individuals often abandon still-living prey after consuming part, usually the head (Fig. 6A, Supplemental Video 3). Worms also sometimes share meals (Fig. 6B,C; Supplemental Video 4). Over many observations in bowls where confinement does not preclude it, satiated worms commenced to swim slowly for some time after feeding, anterior up in the water column and slowly spinning along their axis.

### 3.3 Veliger-eating pelagic flatworms

While the most common type of pelagic flatworm in our collections fed preferentially on nauplii, in late winter we retrieved a very similar type with an apparent preference for molluscan veligers. This was detected as before by keeping worms retrieved from plankton samples in a common bowl with many possible prey, then observing continuously for several hours at a stretch to determine directly who ate whom. The veliger-eaters were somewhat distinct in color (greener) and arrangement of marginal cirri: instead of two closely-spaced posterior cirri embedded within a group of stationary cilia, these had two widely-spaced solitary cirri; they also had at least one pair of lateral “brushes” of stationary cilia with single cirri emplaced within them (Fig. 7). Otherwise these had similar configuration of eyespots, mouth, rhabdites, and digestive system. They attached to gastropod veligers, somehow inserted the pharynx past the operculum, then scooped the prey whole from its shell and swallowed it (Figs. 8). This is a distinct mode of feeding from the nauplius-eaters. These worms also fed on bivalve veligers. They were offered barnacle nauplii and other crustaceans as well, but did not eat them.

**Figure 7.**
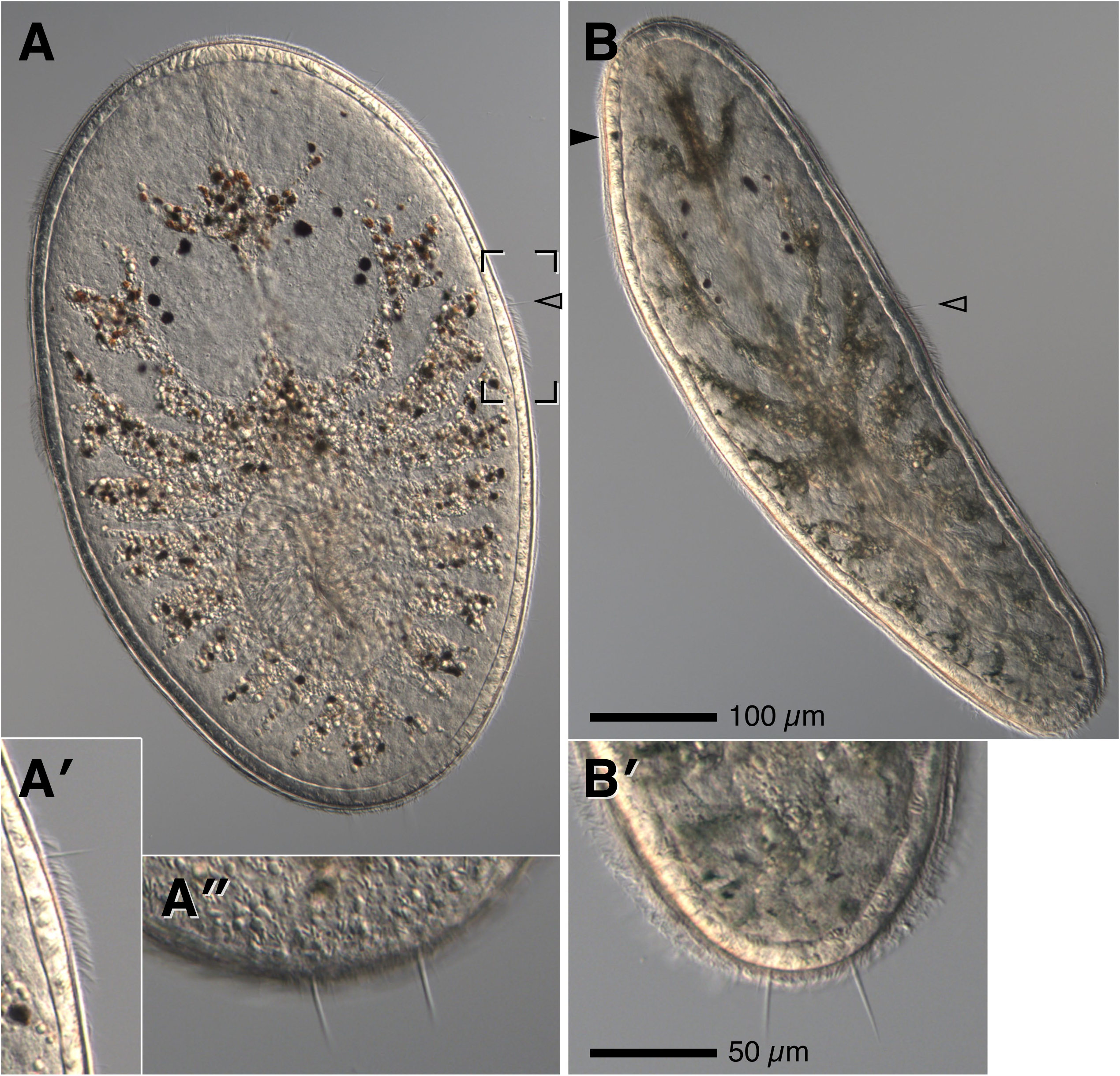
Veliger-eating pelagic flatworms, first morphotype. (A) Spread individual showing similar proportions and arrangement to animals in Figs. 1–3, but with distinct single lateral cirri (hollow arrowhead; also in B) embedded within a batch of stationary cilia (inset A’) and with two widely-spaced discrete posterior cirri (inset A”, different focal plane of the same animal). (B) A swimming individual with greenish tinge and showing a left-side anterior marginal spot (solid arrowhead), likely derived from a larval eyespot. Inset B” is a different focal plane from the same individual; all veliger-eaters of this type had two widely-spaced posterior cirri unaccompanied by other stationary cilia.

**Figure 8.**
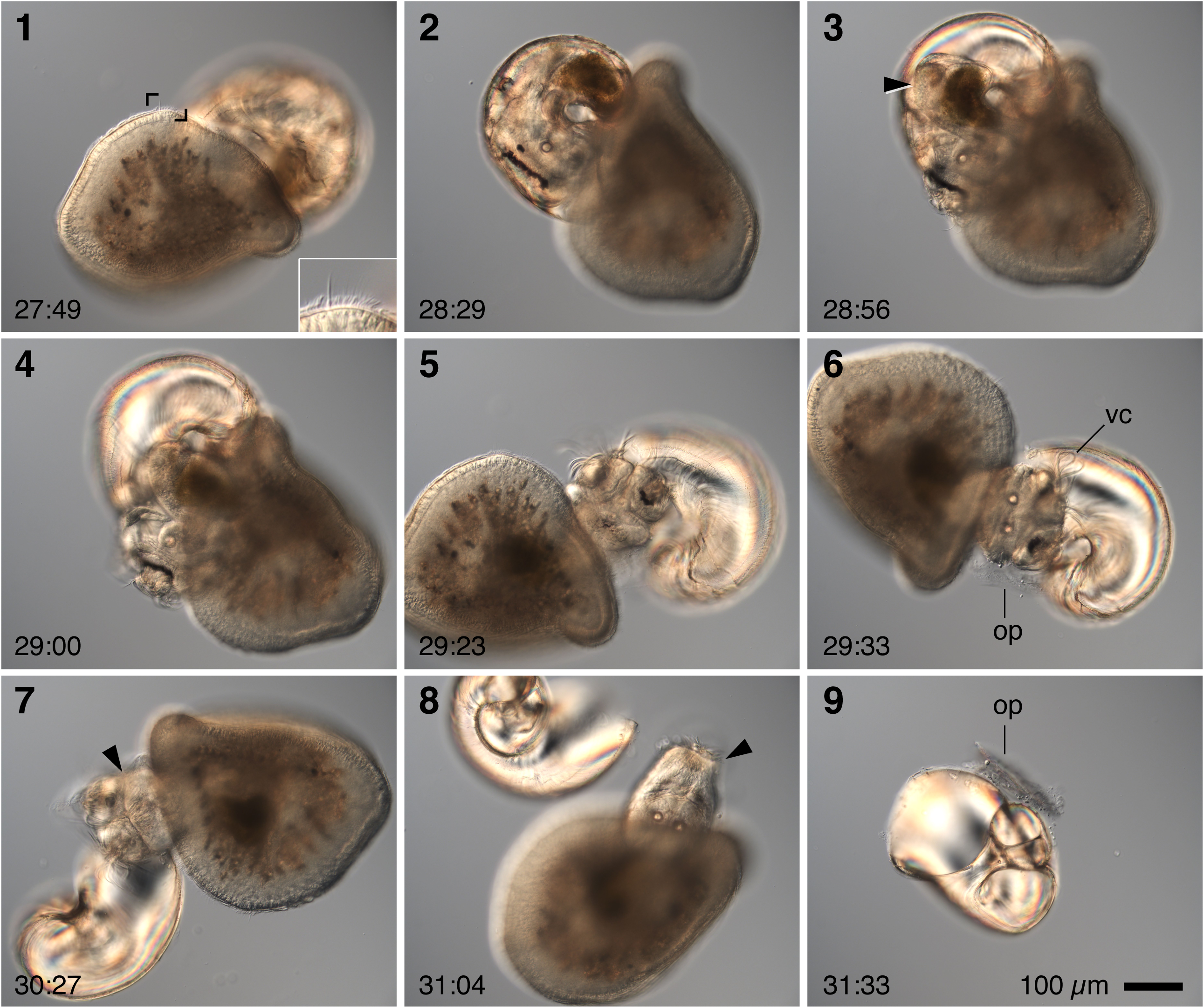
Pelagic flatworm devours a gastropod veliger. Sequence of still images; timestamps are clock time in minutes:seconds. This worm was initially observed swimming near the water surface in a bowl, attached to the already-closed veliger. After they sank to the bottom of the bowl they were transferred together to a slide and carefully coverslipped. Inset in frame 1 is a 3x blow-up showing that this worm has the lateral cirrus-within-brush as in Fig. 7. The worm’s everted pharynx is initially attached to the operculum. In frame 2 the veliger was seemingly secure in its shell, but abruptly began to be pulled forth; in frame 3, the solid arrowhead points out the worms pharynx, intruded all the way around the visceral mass of the veliger, as it pulls the snail from its shell. Perhaps because of the mechanism of scooping out the snail, the flatworm swallowed the visceral mass first, leaving the head and velum for last (frames 4–7; vc=velar cilia). This was observed in nearly all cases. The still-everted pharynx enveloped the head and velum (frames 5–8), separating the operculum (op in frames 6 and 9) and leaving an empty shell.

Further collection yielded a third much more distinct type – larger, thicker, blond, with numerous submarginal spots, and a single posterior cirrus – which also fed on veligers (Fig. 9). Yet later we found three individuals (and more in subsequent collections) that matched the cirrus arrangement of the red-cheeked nauplius-eaters – regularly-spaced anterior cirri, tightly-spaced lateral and posterior pairs – but which were first observed preying upon veligers (Fig. 10 and Supplemental Video 6). Two of these were cultured in isolation with a choice of prey, and each demonstrably ate both barnacle nauplii and gastropod veligers.

**Figure 9.**
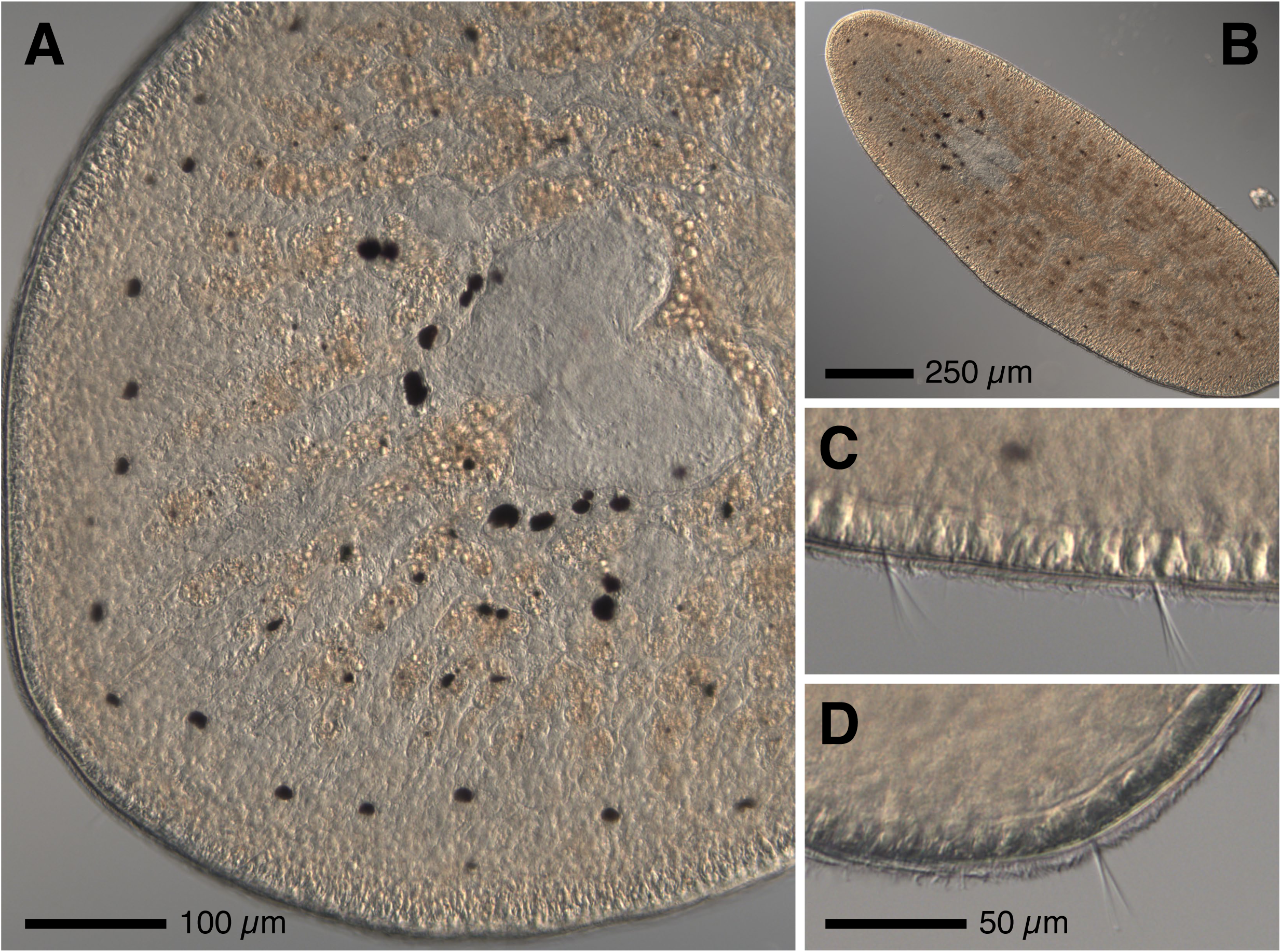
A second type of veliger-eating pelagic flatworm. (A,B) This large individual was captured at >500 µm ECD and has grown in the lab on a diet of wild-caught gastropod and bivalve veligers. The row of regularly-spaced submarginal spots is most prominent anteriorly, but extends all the way around to the posterior (B). The two pairs of outlying eyespots are more superficial and held on stubby tentacles when the animal is unconfined. Animals of this type had several lateral cirri (two adjacent ones shown in C, which is a 2x blow-up of the same individual), usually frayed and not embedded amongst stationary cilia, and a single posterior cirrus (in this case, also frayed: D is a 2x blow-up from the same animal).

**Figure 10.**
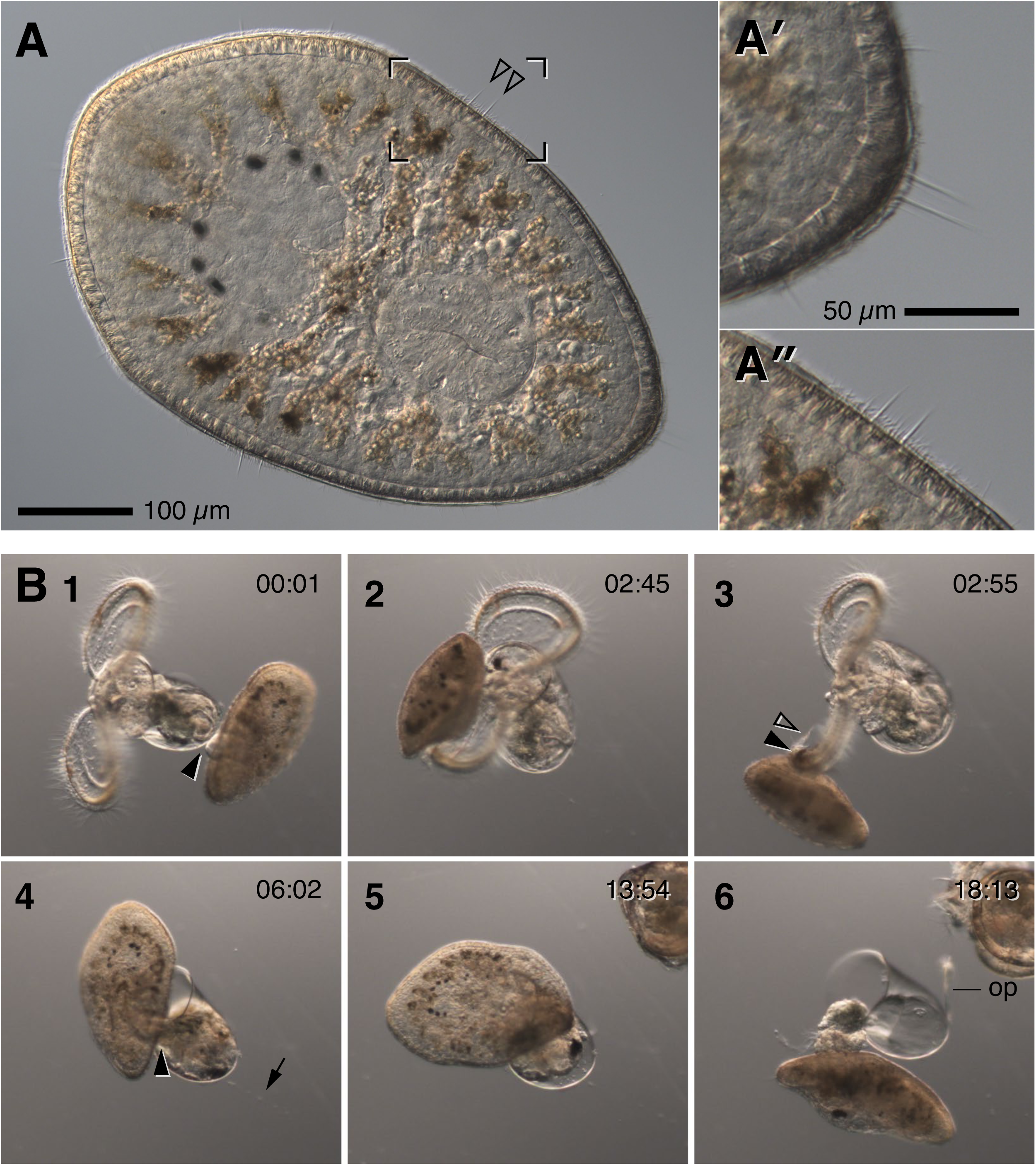
A pelagic flatworm that eats both veligers and crustaceans. (A) This type has two closely-spaced posterior cirri surrounded by long stationary cilia (inset A’ is a 2x blow-up from a different focal plane of the same individual) and flanked by two shorter cirri, and closely-paired lateral cirri (inset A” from the region shown by brackets in A). (B) Frames from a real-time video sequence in which the same individual in (A) devours a large gastropod veliger; corresponds to Supplemental Video 6. Initially the worm attached by its everted pharynx (solid arrowhead, frame 1) to the shell opposite the aperture. It spent many minutes thus attached, as the gastropod extended its velum and beat vigorously. However, the flatworm appears to have tethered the veliger to the substrate: a mucus strand is faintly visible frames 2–5 (arrow in frame 4, where it is clearest). At some point the worm detached from the shell, and crawled over the velum (frame 2) which the snail inexplicably failed to retract; then the worm grabbed the velar margin with its pharynx (solid arrowhead, frame 3), apparently applying mucus in the process (hollow arrowhead, frame 3), whereupon the snail shortly thereafter withdrew into its shell. The worm intruded its pharynx past the operculum (solid arrowhead in frame 4), and ultimately scooped out the snail, devouring it entirely while leaving behind empty shell and detached operculum (op in frame 6).

The worms we captured in this study were offered a diverse menu including not only crustaceans and bivalves, but also rotifers, various annelids, actinotrochs, hoplo- and palaeonemertean larvae, and various ciliates. We do not claim to have sampled exhaustively nor offered a full spectrum of possible prey, but aside from occasional encounters – one observed act of cannibalism and another instance in which a small trapped spionid was devoured – we have yet to find any pelagic flatworm of this general type that regularly preys upon any other animals besides planktonic crustaceans and molluscan veligers.

### 3.4 Growth in culture

The general spectrum of sizes at capture suggests these are hatchlings which grow during a planktonic phase. We measured four separate collections from two widely-separated times (late March and early May): to minimize variation in shape and posture, worms were slightly flattened under a coverslip to as close to standard thickness as possible (restrained but still able to move), then photographed; each worm’s area was measured, then twice the square root of area/π taken as an equivalent circular diameter (E.C.D.). Out of 43 animals, about a third (15) fell into the 150-250 micron bin (the smallest in these collections was 168 µm E.C.D.), another third (16) in the 250-350 micron bin, and the rest larger (Fig. 11A; the two largest in this subset were 459 and 551 µm E.C.D.).

**Figure 11.**
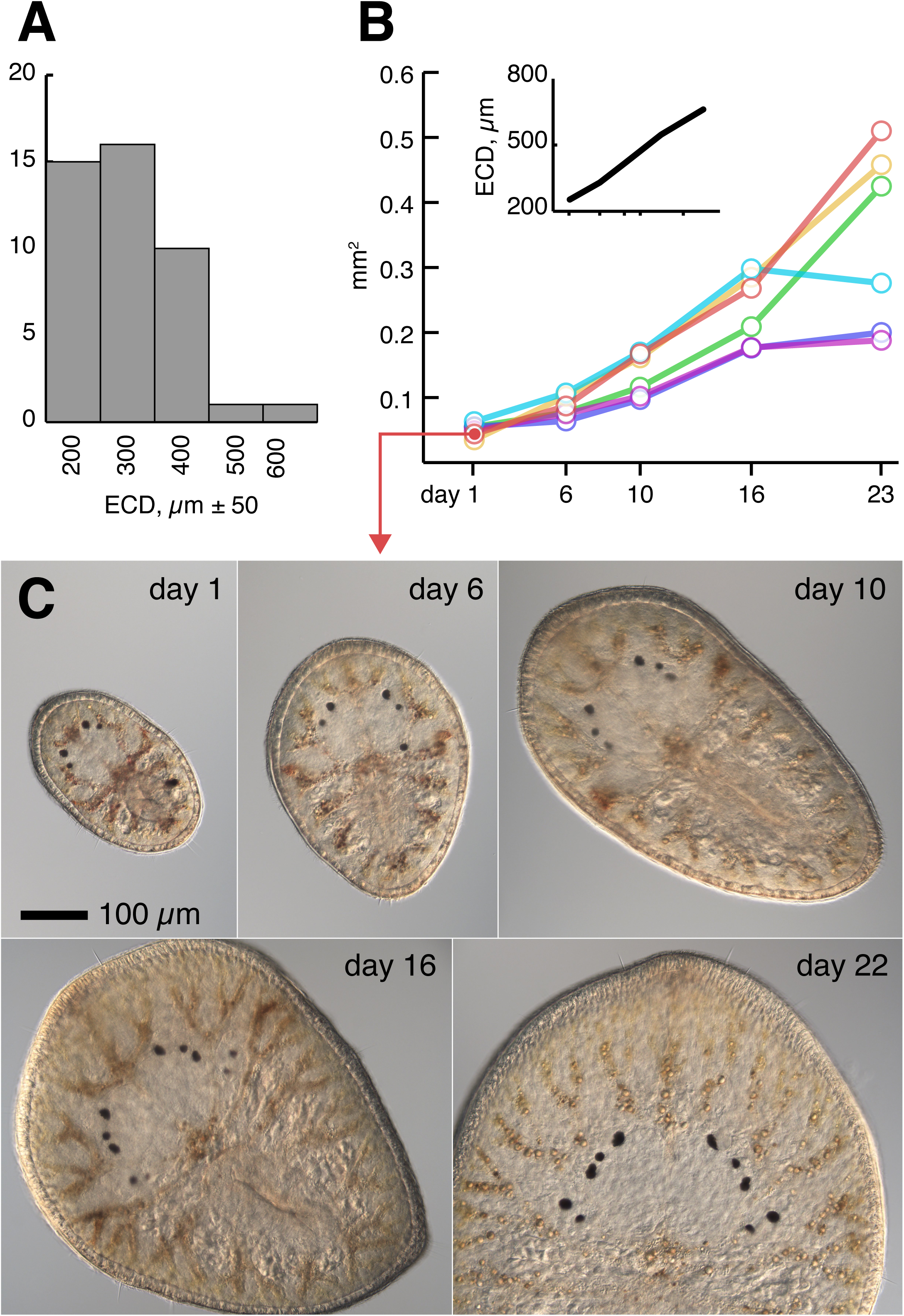
Pelagic flatworms grow substantially on a carnivorous ration. (A) Histogram of size at capture, before feeding in lab. Area of flattened wild-caught larvae was measured from calibrated images and equivalent circular diameter (ECD) calculated. Numbers represent four pooled collections separated by six weeks. (B) Three-week growth of six isolated individuals fed barnacle nauplii. Inset shows average ECD for the same set. (C) One of the individuals from (B), at each measured timepoint.

Wild-caught animals grew in communal bowls, but at uneven rates – possible reasons include unnoticed collection damage, competition for prey, variation in prey quality, disease in captivity, or undetected taxonomic variation. Therefore for a rough estimate of growth rate, we isolated nine small individuals of the pale nauplius-eater morphotype from two successive day’s collections into separate bowls and provided each with a dozen or so newly-released *B. glandula* nauplii. These were photographed and measured regularly over three weeks, with refreshment of water and prey (the nine initial measurements from these animals are included above). Three were lost. The remaining six grew steadily, on average quadrupling in size over 16 days (Fig. 11B,C). At about the time they reached ∼1 mm long, these, and others in group captivity, ceased to swim and became sticky and difficult to transfer. They also became less and less inclined to attack and devour now-much-smaller barnacle nauplii, and began to shrink instead of grow. These gliding, sticky flatworms we provisionally regard as true juveniles.

Juveniles were easily lost in culture, as they developed a tendency to crawl beyond the water surface, thereafter drying on the side of the bowl. The few survivors were offered the smallest barnacles available, obtained live in groups on small rock chips chiseled from nearby shores. Upon feeding on these, the juveniles darkened markedly in color and recommenced growth (Fig. 12). While the pelagic larvae may have been positively phototactic, the true juveniles tended to move away from light sources and generally hid under rock chips or in empty barnacle tests. Feeding was observed by allowing worms to initiate meals in dim light. Feeding also involved behaviors somewhat distinct from the mode of the larvae: the juveniles moved from barnacle to barnacle, seeming to test them somehow, before choosing a victim. They always chose live victims, but avoided ones combing vigorously. Having chosen a meal, worms inserted their posterior end into the top to cover the opercular plates completely, and seemingly waited for some opportunity for as much as an hour. Worms sometimes moved off to another victim, or fled the scene entirely after an injudicious adjustment of the light. At some point the worm everted its pharynx either between the operculum and the marginal plates, or through the aperture, and soon thereafter began to pump barnacle tissue into its digestive diverticulae. Ingestion took tens of minutes.

**Figure 12.**
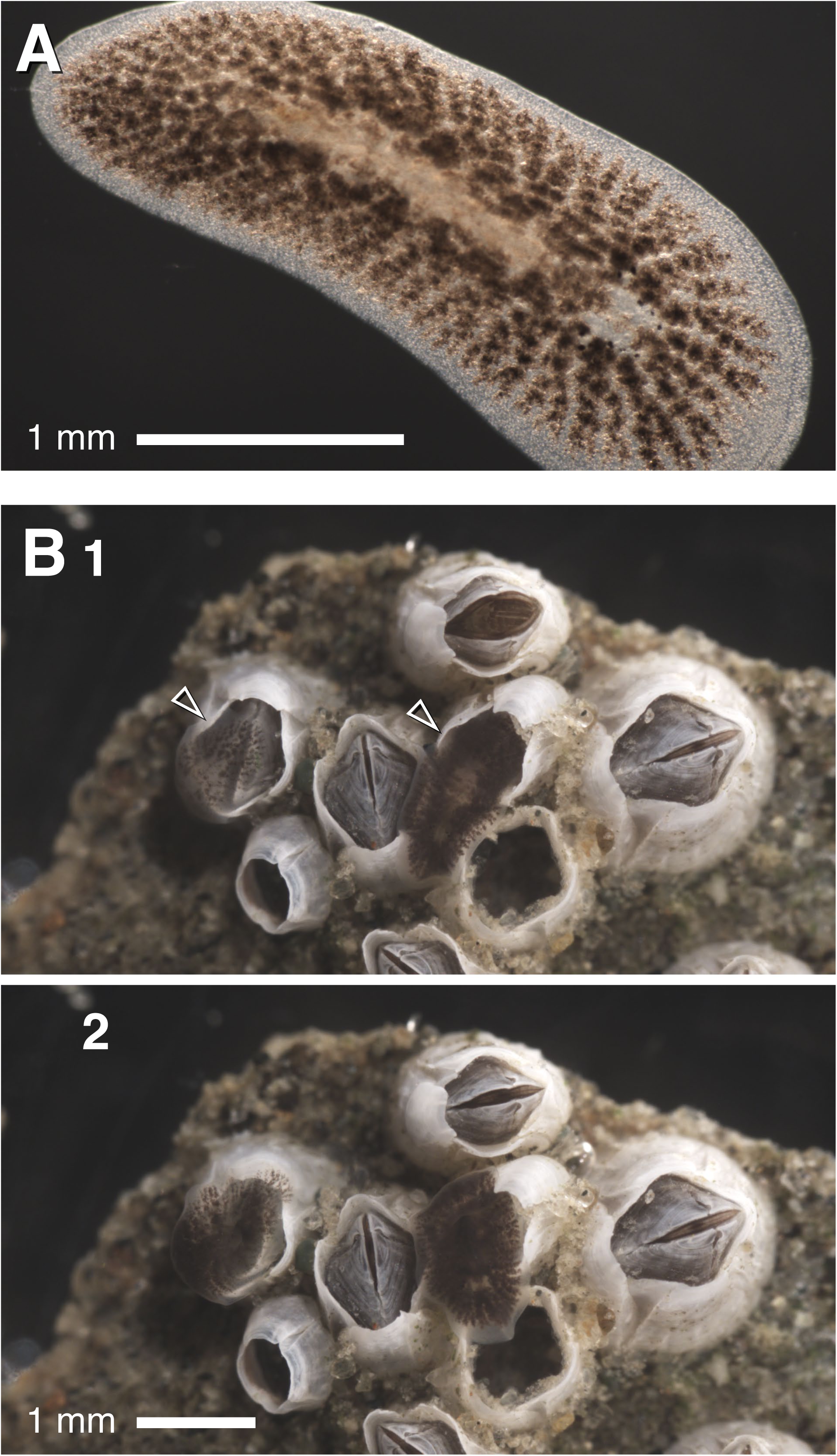
Juvenile flatworms of the nauplius-eating type. (A) This animal was once the type shown in Fig. 1; it has darkened markedly after consuming small barnacles, sticks to surfaces on which it glides, and no longer swims with ciliary motion. (B) Two juveniles (arrowheads) feeding on small *Balanus glandula*. In the first frame, both rest their posterior portion across the operculum. In the second, both have inserted their pharynx and have darkened further as they pump prey tissue into their digestive system. The left one inserted through the aperture; the right one inserted between operculum and marginal plates.

Two of these individuals, cohabiting a single bowl, grew to >1 cm body length within two months of commencing to feed upon adult barnacles. Thereupon they began to lay eggs. These were pale and yolky and averaged 140 µm in diameter (±8.6 µm; n=55 from two different egg plaques), deposited in tightly-packed monolayers on the glass. Many initiated development; after three weeks, embryos slowly rotated within their capsules but had no visible anatomical features, only irregular globular masses beneath a thin ciliated epidermis. In the fourth week, a pair of eyespots became apparent and some muscular contraction was detected. Precisely one month after the first egg mass was found, at ambient temperature of 14-16C, hatchlings began to emerge. These were keyhole-shaped with two large and two small eyespots, as Fig. 1C, and 150-200 µm (158±11 µm in E.C.D., n=19). Curiously, many had five or even six eyespots upon emergence and some irregularity in the position of the posterior cirri, but within a day or two most had only four eyespots and two closely-spaced posterior cirri.

These hatchlings initiated feeding within hours of emergence from egg capsules. They were offered nauplii of *B. glandula*, whereupon – as their parents had done when wild-caught from plankton – each hatchling selected one, circled around it, and attached using everted pharynx, usually to the dorsal carapace of the prey. It is particularly noteworthy that the 1st and 2nd instar nauplii are nearly an order of magnitude larger than the flatworm hatchlings (272±9 µm long, not including appendages or caudal spine, n=25; the nauplii are thicker than the flatworms and appendages represent a significant fraction of tissue). Comically, the still-swimming nauplius appeared to be wearing a large beret; but it appeared that the hatchlings must entangle the nauplii with invisible mucus, because soon they became stuck to the glass of the bowl or to the surface film, whereafter they were devoured, often by several hatchlings at once who recruited successively to the captured prey. Some hatchlings grew rapidly, but a large fraction did not; we do not know whether this was due to sub-optimal culture conditions or intrinsic challenges of macrophagous feeding.

### 3.5 Hatchlings of *Notocomplana litoricola* are planktonic predators

Mid-spring, we began to catch small flatworms similar to the nauplius-eaters, but darkly pigmented purplish-brown throughout the epidermis (Fig. 13A,B). These matched the nauplius-eaters in cirral arrangement, and also attacked and devoured nauplii eagerly. These were reminiscent of hatchling “juveniles” of a locally-abundant polyclad, tentatively identified as *Notocomplana litoricola* HEATH & MACGREGOR 1912 (following description in Hyman, 1953; and keys by Holleman, 1987 and 2007). This animal as an adult is mottled tan to dark gray or brown, 1-4 cm with an axial ratio of 2–4, lacks tentacles, is found amongst and feeds on high-intertidal barnacles, and swims actively when disturbed (our obs.). (Shinn, 1987, also describes similar “black” hatchlings of *Notocomplana* (=*Freemania*) *litoricola*, adults of which he says accept limpets and mussels as food; our collected worms ate only barnacles.). We therefore acquired several adult *N. litoricola* from undersides of high intertidal rocks in the Coos Bay estuary (Fossil Point) and outer coast (Lighthouse Beach) and brought them to the lab, whereafter they promptly laid numerous egg plaques. These developed over the course of approximately four weeks to hatch into swimming flatworms identical in every respect with the dark nauplius-eaters recovered from plankton samples (Fig. 13C). Although when laid the eggs were blond, and remained so during most of embryogenesis, they darkened abruptly as they approached hatching and the hatchlings were nearly black. Hatchlings of *N. litoricola* were larger (218±14 µm E.C.D., n=19) than either the lab-reared hatchlings of the pale nauplius-eaters, or the smallest wild-caught larvae. When offered barnacle nauplii, the hatchlings promptly latched onto and devoured them, growing rapidly (Fig. 13D; 330±39 µm E.C.D, n=15 after 4 days’ feeding, likely corresponding to ∼3-fold increase in mass; 376±28 µm E.C.D., n=12 after 6 days, ∼5-fold increase). They also consumed decapod zoeas (Fig. 6C and Supplemental Video 5) and copepods (not shown).

**Figure 13.**
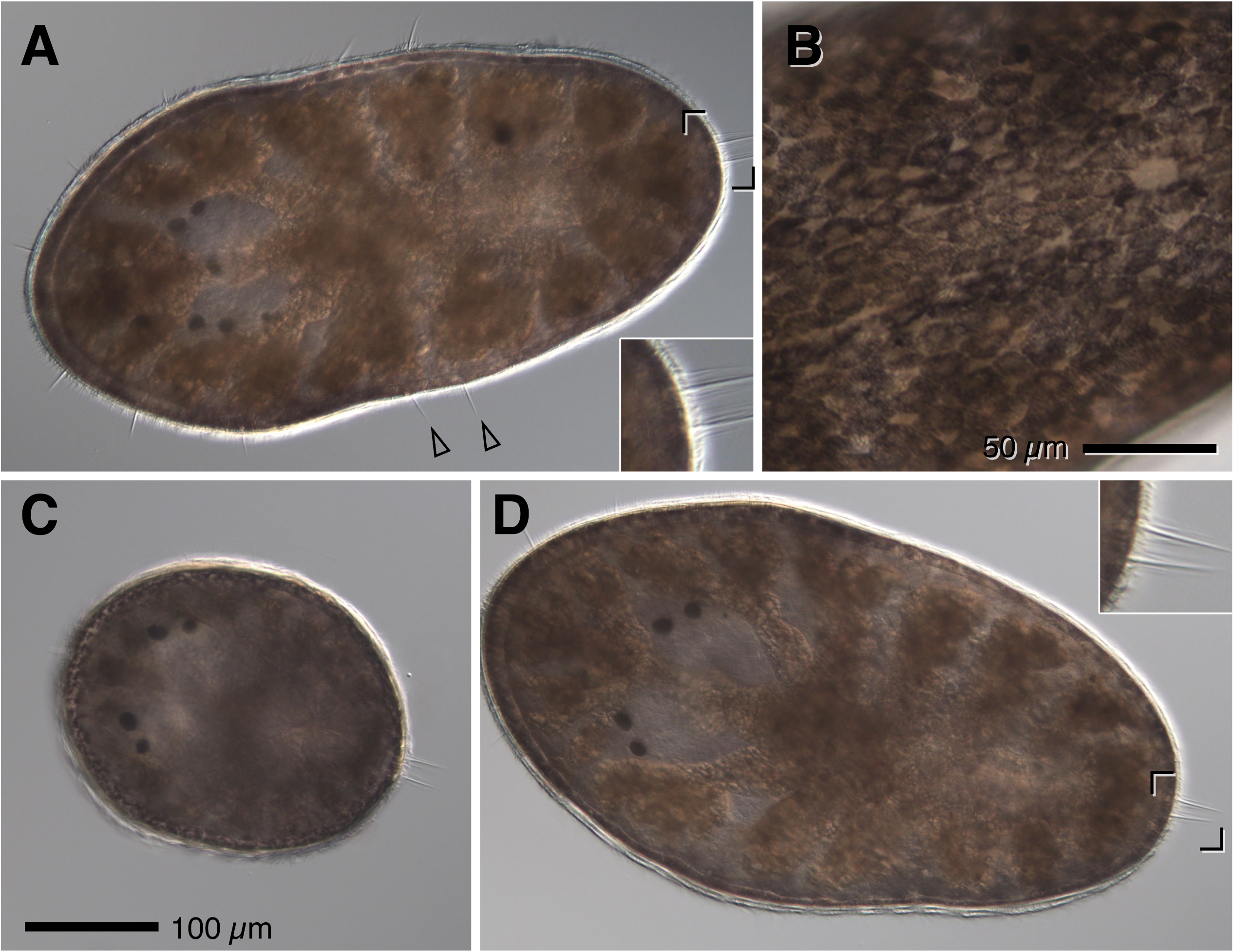
Dark nauplius-eaters are hatchlings of *Notocomplana litoricola*. (A) Wild-caught dark nauplius eater with arrangement of cirri and proportions identical to the morphotype in Fig. 1. Inset is a 2x blow-up showing posterior cirri, surrounded by long single non-motile cilia; hollow arrowheads indicate the two well-separated lateral cirri. There are also six short and well-separated anterior cirri, as in all other morphotypes considered here. (B) Superficial focus on the same individual: all the dark pigment appears to be due to epidermal cells. (C) Newly-emerged hatchling from collected *N. litoricola*, before feeding. Note posterior cirri and two pairs of cerebral eyespots. (D) An individual from the same cohort as (C), after two days’ feeding on barnacle nauplii. Inset is a 2x blow-up of posterior cirri for comparison to (A).

### 3.6 Some Müller’s larvae metamorphose into swimming carnivores

One of us previously studied feeding on phytoflagellates by wild-caught Müller’s/Götte’s larvae of several unknown species of polyclad (von Dassow & Ellison, 2020). The most abundant of these is a stout type with four to six short lobes and a mix of black, brown, green, and rust-red endodermal pigmentation (Fig. 14A; the “Winnie-the-Pooh” morphotype *sensu* von Dassow & Ellison, 2020) which DNA barcoding broke into two OTUs. These larvae, in lab culture, never progress beyond six lobes and ∼250 µm, and metamorphose into an atentaculate ovate form that swims continuously (as opposed to crawling on a surface; Fig. 14B,C). These metamorphosed Müller’s larvae are similar in size, color, and other features to the small pelagic flatworms we caught in this study. In fact, we initially began to keep these small worms from plankton tows because we suspected they might be the juveniles of the Winnie-the-Pooh larval morphotype.

**Figure 14.**
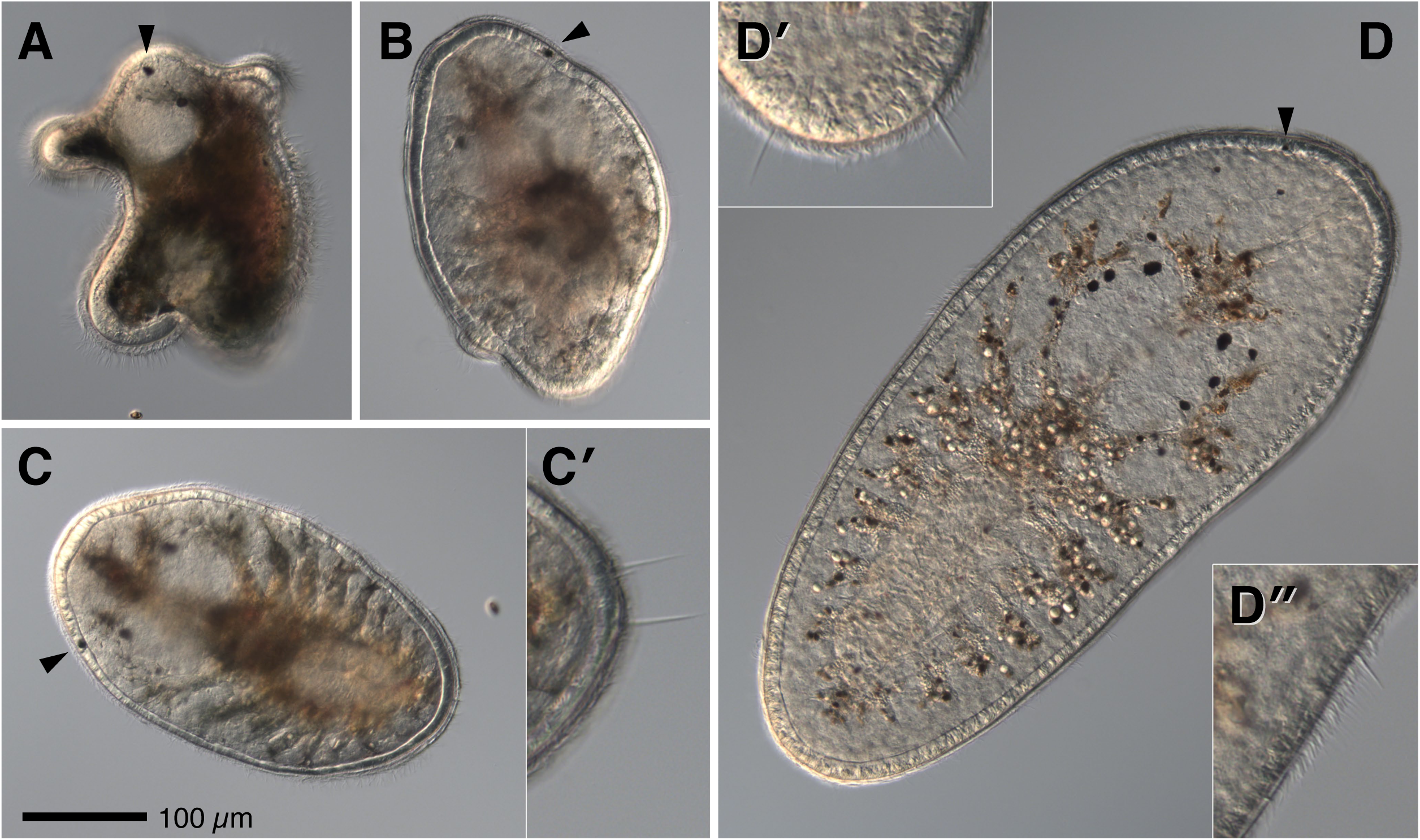
One kind of Müller’s larva transforms into a veliger-eater. (A) Wild-caught Winnie-the-Pooh type Müller’s larva that has been feeding in lab culture on *Rhodomonas lens*. This focal plane includes an anterior left-side superficial eyespot (arrowhead). (B) Partly-metamorphosed Müller’s larva, from the same collection as (A), with remains of heavily ciliated oral lobes still present, and no clear pharynx yet. Arrowhead indicates left-side superficial eyespot (this is an oral view). (C) Fully-metamorphosed Müller’s larva, possibly same individual as A. Pharynx has formed, and this animal was observed devouring a veliger shortly after this photo was taken. Arrowhead indicates left-side superficial eyespot. Inset (C’) is a 2x blow-up of a different focal plane of the same individual to show the posterior cirri, well-separated and not accompanied by stationary cilia. (D) A former Müller’s larva after feeding for several weeks on small wild-caught veligers. This animal was still a swimmer. Note left-side eyespot (arrowhead). Insets (D’ and D”) are 2x blow-ups from different images of the same individual to illustrate posterior and lateral cirri.

Once we discovered that small planktonic juvenile flatworms eat large prey, we wondered whether metamorphosed Müller’s larvae might adopt similar habits. In 2020, Müller’s larvae were not numerous in our plankton samples. We retained approximately a dozen of the Winnie-the-Pooh morphotype over the course of several tows. These were cultured together in a single bowl with ample supply of *Rhodomonas lens* for food. Gradually over the course of several weeks these were either lost or underwent metamorphosis. At this point we added a few dozen barnacle nauplii or wild-caught small veligers to the culture bowl. Examining the bowl daily revealed one or two newly empty veliger shells, but no apparent losses of nauplii due to predation. We ceased adding nauplii and provided more gastropod and bivalve veligers, both of which we subsequently found empty shells of. Out of four surviving juvenile worms, three clearly grew larger (Fig. 14D), and finally each was observed devouring a gastropod veliger at least once (empty shells indicated that some had eaten bivalves as well). After several weeks in this state, these individuals still swam continuously rather than gliding on surfaces.

From a subsequent collection of four such Müller’s larvae, three promptly metamorphosed after several days, swimming continuously and exhibiting the same cirrus arrangement as the previous set, whereupon they too were offered gastropod veligers from plankton in a small Syracuse dish. Within minutes of adding prey, all three attached to veligers larger than their own size, dragging them down to the bottom of the dish. Observing continuously over the course of the next hour, each successfully scooped the entire body of the veliger from its shell and engulfed it velum last, as we observed for the wild-caught worms.

### 3.7 DNA barcoding confirms that discrete morphotypes are likely distinct species

We isolated individual wild-caught planktonic worms for DNA extraction after observing them eat at least once. Individuals were kept in clean seawater for one day after a meal, then rinsed in filtered seawater and frozen. Results are summarized graphically in Fig. 15. The majority of specimens yielded sequences for both 16S and COI (accession numbers in Table 1), although despite one day’s fasting, several of the recovered COI sequences represented an apparent meal.

**Figure 15.**
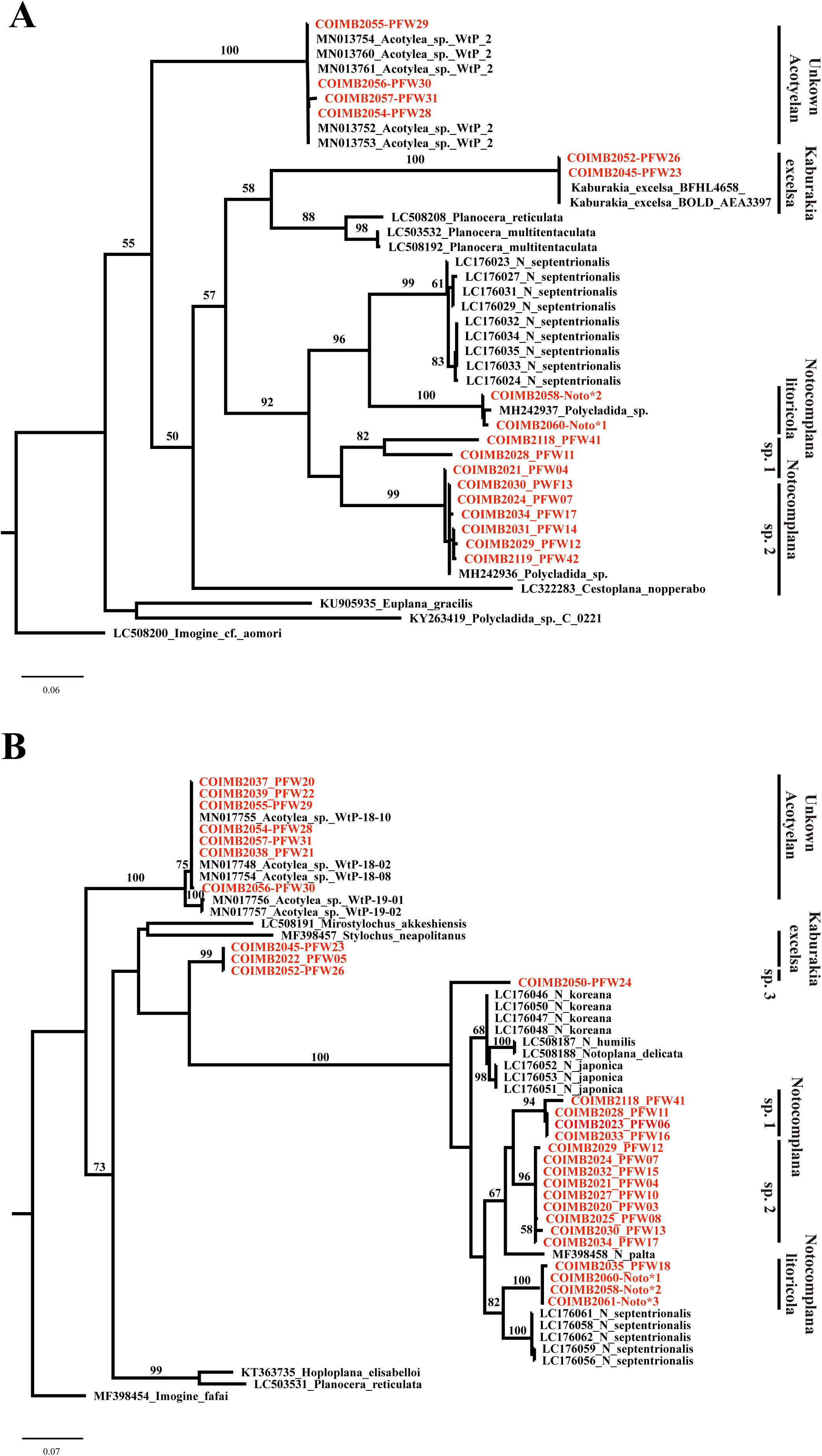
Resulting trees from the Maximum Likelihood analysis with RAxML from pelagic flatworm samples and GenBank sequences. Support values above 50 on each branch A: COI phylogeny (lnL = -4504.5267). B: 16S rRNA phylogeny (lnL = -2293.6066). Samples in black from GenBank. Vertical blue bars indicate groups delimited by ABGD. See Table 1.

Sequencing revealed, first of all, that 14 sequenced nauplius-eaters are related (but not identical) to already-sequenced species of *Notocomplana* (16S and CO1 both ∼95% identical to *N. palta*). Of these, 10 constituted one OTU (corresponding to the morphotype in Fig. 1; *Notocomplana* sp. 2) and four, which shared extensive patterned red pigmentation (Fig. 2; *Notocomplana* sp. 1), formed another. Surprisingly, the visually-distinctive type characterized by large orbs in the digestive diverticula, with red and yellow pigment patches in the epidermis (Fig. 3), belonged to the dominant nauplius-eater group (*Notocomplana* sp. 2). Before sequencing, we had determined that this morphotype remains distinctive for at least several days in fed culture, hence it seems unlikely to be diet-dependent.

Second, a single dark-pigmented wild-caught nauplius-eater was successfully barcoded and matched sequences from three locally-collected adult *Notocomplana litoricola* (neither *N. litoricola* or other local species in the genus is yet represented in Genbank; the closest publicly-available sequence is ∼95% match to *N. septentrionalis*). These formed a third OTU, distinct from the other nauplius-eaters. Only one of the “generalist” type that ate both veligers and nauplii (Fig. 10) could be successfully barcoded, but that one 16S sequence nevertheless placed it outside the other *Notocomplana* OTUs (*Notocomplana* sp. 3).

A fifth OTU lacked close similarity to other previously-sequenced polyclads, and included two large veliger-eating specimens with marginal spots (Fig. 9) as well as a single smaller, red-tinged specimen that appeared similar in general type to others but with distinct swimming behavior and cirrus arrangement. Subsequently we were informed by G. Paulay that these sequences closely match *Kaburakia excelsa* BOCK, 1925 collected from San Juan Island, WA.

Finally, a sixth group included three individuals from the group of veliger-eaters with widely-spaced posterior cirri (Fig. 7), which matched 100% with sequences we retrieved (and deposited previously; von Dassow and Ellison, 2020) from Winnie-the-Pooh type Müller’s larvae, which are the offspring of an unknown acotylean. These also matched three sequenced individuals that we newly caught in plankton samples as Müller’s larvae, which grew on *Rhodomonas* in culture, then transformed into veliger-eating swimming wormlets (Fig. 14).

These six groups are recovered in phylogenetic and ABGD analysis. ABGD results for COI sequences found a barcoding gap at K2P distance of 0.001–0.060, and for 16S rRNA sequences a K2P distance of 0.0028–0.036. Both fragments recovered the same six groups seen in the phylogenetic trees (Fig. 15).

### 3.8 DNA barcoding identifies prey remains in planktonic polyclads

Despite care to deprive captive wild-caught polyclads of food for at least a day prior to preservation for DNA barcoding, some individuals yielded COI sequences matching crustaceans that had been offered in the lab as prey (data not shown). This encouraged us to seek evidence for predation in nature by a similar approach. Because sampling unavoidably concentrates the predators with their potential prey, it is always possible that the gut contents of any individual flatworm retrieved from a plankton sample reflect encounters within sample confines rather than in nature. To limit this possibility, we reasoned as follows: most acts of predation recorded in the lab take place over the course of ∼30 min.; initial stages of predation are easily disrupted, i.e. worms can be knocked off selected prey easily; once a meal has commenced, the worm and its prey generally sink to the bottom; finally, in a fresh plankton sample, swimming worms collect on the lit side of a jar, near the surface, where also swarm myriad active crustaceans, under which circumstance the worms appear unable to select and attach to prey due to incessant battering. We therefore collected, on several successive days, as many polyclads as possible from the lit-side surface, within the first 30 min. after collection. These individuals were isolated immediately into clean filtered seawater. Those with apparent gut contents were then photographed, rinsed in clean seawater, and frozen. 10 out of 24 specimens so prepared yielded single COI sequences matching possible prey (Table 2, Figure 16): six near-perfect matches to various barnacles (*Balanus glandula, Balanus crenatus, Chthamalus dalli*; all locally abundant), two near-perfect matches to a bivalve (*Adula californiensis*), and two poor matches to a rotifer (*Synchaeta* sp.). These results are graphically summarized in Figure 16.

**Table 2.** Specimens of pelagic flatworms that yielded apparent prey sequences. Corresponds to Fig. 16. Predator ID is based on microphotographs showing arrangement of marginal cirri.

| Specimen ID | Predator ID (Based on morphology) | Closest GenBank Match (Percentage of similarity) | GenBank Accession COI |
| --- | --- | --- | --- |
| COIMB2078 | Notocomplana sp. 3 | EF687902 - Balanus glandula | MW256479 |
| COIMB2079 | Notocomplana sp. 3 | EF552057 - Balanus glandula | MW256481 |
| COIMB2083 | Notocomplana sp. 3 | EF552057 - Balanus glandula | MW256480 |
| COIMB2084 | Notocomplana sp. 3 | MG422323 - Adula californiensis | MW256487 |
| COIMB2085 | Notocomplana sp. 3 | AF234814 - Chthamalus dalli | MW256484 |
| COIMB2086 | Notocomplana sp. 3 | MG421601 - Adula californiensis | MW256488 |
| COIMB2103 | Notocomplana sp. 2 | KY235399 - Balanus crenatus | MW256482 |
| COIMB2105 | Notocomplana sp. 2 | MT736465 - Balanus crenatus | MW256483 |
| COIMB2107 | Notocomplana litoricola | LC215583 - Synchaeta sp. | MW256485 |
| COIMB2108 | Notocomplana litoricola | LC215583 - Synchaeta sp. | MW256486 |

**Figure 16.**
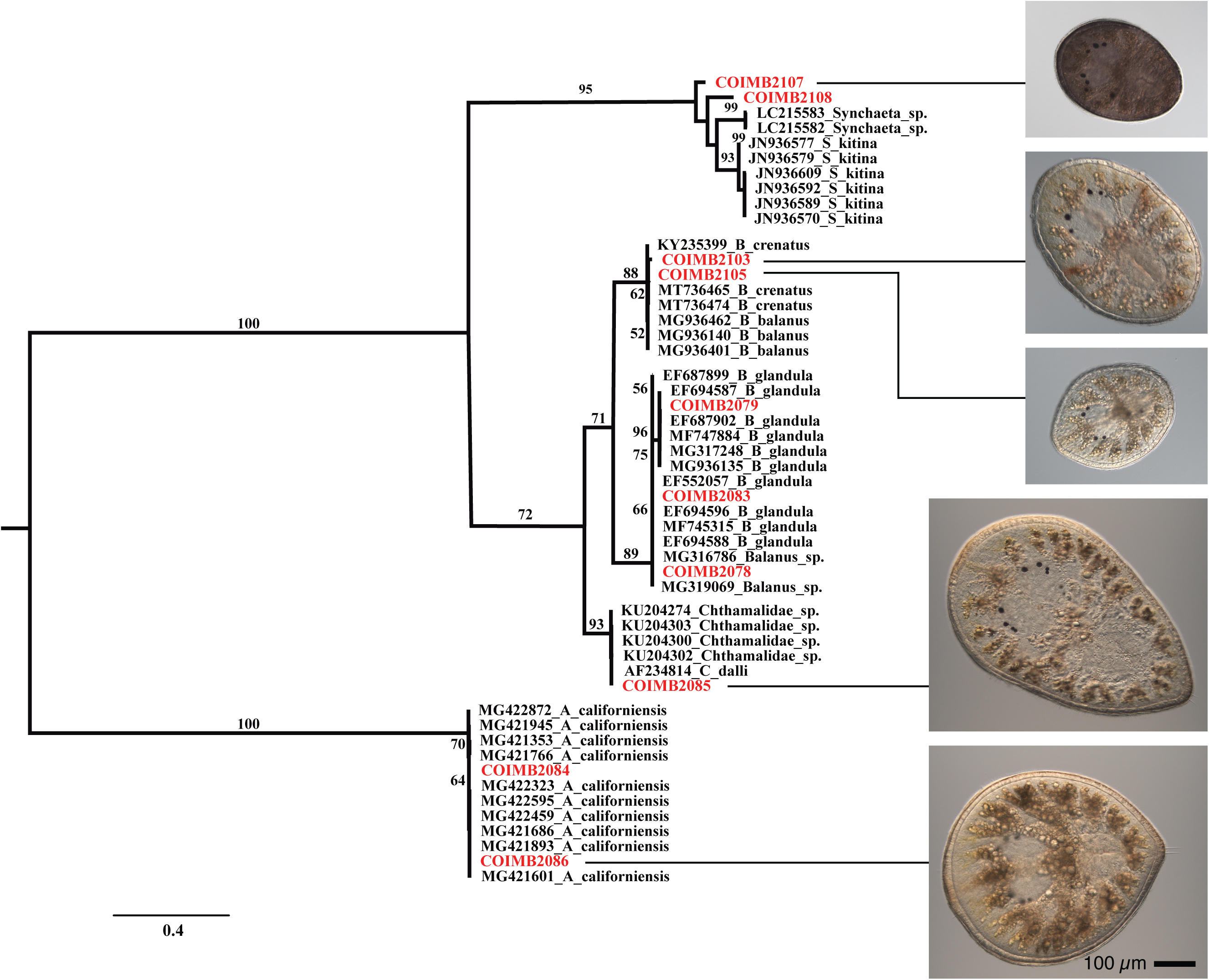
Resulting tree from the Maximum Likelihood analysis of COI sequences (InL = -3780.8591) from food content of pelagic flatworm samples and GenBank sequences. Support values above 50 on each branch. Samples in black from GenBank. Names in red indicate the samples from which the apparent gut contents sequences were extracted. Representative images of specimens tentatively identified by color and cirral arrangement as *Notocomplana litoricola* (COIMB2107), *Notocomplana* sp. 2 (COIMB2103 & COIMB2105; compare to Fig. 1), and *Notocomplana* sp. 3 (COIMB2085 & COIMB2086; compare to Fig. 10)

The recovery of apparent rotifer sequences from these specimens, none of which had any chance to feed in the lab, was a surprise, as we had never observed predation upon rotifers despite offering them routinely as part of the lab menu. Because some predatory copepods likely eat rotifers, and during our attempts many copepods amplified poorly when using universal COI primers, it is conceivable that these rotifer sequences actually represent gut contents of crustaceans that had been preyed upon in turn by flatworms.

## 4. Discussion

We have identified a group of pelagic flatworm juveniles (or larvae) that feed on crustacean or molluscan larvae by macrophagous carnivory – direct interception, penetration, and whole ingestion. Are these really pelagic? We catch them in plankton nets, but of course we also catch spruce needles and other organisms that end up in the water from other habitats. The habit of extended ciliary swimming, however, and the apparent preference for an indubitably pelagic prey seem to affirm that these are indeed pelagic animals. They showed no interest in burying themselves in sediment when offered. In plankton collections, they were recovered almost exclusively from the illuminated side of the jar or bowl, where many other active swimmers also collect due to positive phototaxis, but it was not clear that the worms themselves are phototactic. They may instead follow cues to swim toward their prey, consistent with a pelagic habitat. Even so, it is curious to imagine how these small and relatively slow swimmers encounter their fast-moving prey in open water.

The most noteworthy aspect of our findings is not the mode of feeding or the choice of prey, but that these flatworms are of the size and shape to be considered crawl-away juveniles. In many invertebrate phyla, a stereotypical categorization of developmental modes (e.g., as classically presented by Jägersten, 1972) distinguishes indirect-developing planktotrophs, usually developing from small eggs to graze in the plankton on small prey like phytoplanktonic flagellates, versus direct developers that are generally lecithotrophic and develop from larger eggs, achieving juvenile morphology on maternal investment and adopting more or less the lifestyle and habitat of their parents upon the completion of embryogenesis (with perhaps a dispersal stage or period interposed). An extreme exemplar of this dichotomy – along with associated bimodal distribution of egg size – is provided by echinoderms (Emlet et al., 1987). Polyclad flatworms include many species that develop via a long-lived, planktotrophic Müller’s or Götte’s larva; the rest are considered direct developers (reviewed by Martín-Durán and Eggers, 2012; Rawlinson, 2014), and, in sharp contrast to some other invertebrate groups, there appears to be little evidence for a correlation between egg size and developmental mode among them. The wild-caught pelagic flatworms we studied clearly overlap the size range expected for an invertebrate egg that gives rise to a planktotrophic larva, and the smallest are likely hatchlings based on the known sizes of eggs laid by these and similar species. Yet they are inferred to be feeding in the plankton, living and growing in a mode and habitat distinct from their parents. By this token they are larvae, although by morphology they are juveniles.

Indeed, one type clearly matches hatchlings of *Notocomplana litoricola* in anatomy and behavior. The adult is a benthic crawling predator that consumes barnacles (although it can also swim by undulating its margin), whereas the hatchling is a pelagic ciliary swimmer that consumes barnacle larvae. A recent paper reported laboratory culture of three species of polyclad with “direct” development (including a species of *Notocomplana)* on a diet of cooked brine shrimp (Morita et al., 2018). These authors noted that the hatchlings in their study are swimmers and pointed out the implication for life history and ecology if it were shown that the same animals have an equivalent habit in nature. This we have tentatively shown, albeit for different species and indirectly. In future work we will seek to identify prey remains systematically from natural plankton samples; lab access restrictions this season due to pandemic shutdown limited our efforts. Another related but unresolved question is whether the apparent prey preferences we observed for various morphotypes represent natural specializations or not. At least one type was observed to take both crustaceans and mollusks as prey, but otherwise the worms in our collections showed consistent, nearly exclusive preferences during culture.

A further surprise in our study is that at least some conventional planktotrophs – ciliary swimmers that feed on unicellular phytoflagellates – metamorphose to become pelagic predators on large animal prey, rather than continuing to grow on phytoplankton until they metamorphose and settle as benthic adults. This is unlikely to pertain to all Müller’s larvae, since some are known to achieve large size with elaborate tentacle arrangement (Rawlinson, 2014), but it raises several interesting possibilities. First, it implies that these species have (at least) a three-phase life cycle: first as conventional planktotrophic larvae, then as macrophagous planktotrophs, then as benthic predators. A second larval stage is well-known in a few invertebrate groups – for instance, the barnacle cyprid or the holothuroid doliolaria – but these are post-growth non-feeding forms associated with metamorphosis and site selection.The closest analogous case we can think of are decapods, in which the megalopa or other post-larva succeeds the zoea as a morphologically distinct larval form with a different swimming and feeding mode. It prompts us to wonder whether other planktotrophic larvae also switch feeding mode, with or without morphological change, as they grow large.

Second, the observation that a Müller’s-type larva becomes a planktonic carnivore before settlement raises the possibility that such a switch in mode and morphology could be both phenotypically and phylogenetically plastic. Götte’s larva has sometimes been suggested to be a reduced, lecithotrophic vestige of Müller’s larva (Anderson, 1977; Ruppert, 1978), and was only recently shown definitively to feed (Allen et al., 2017; von Dassow and Ellison, 2020). There are known cases in which closely-related species hatch as feeding versus non-feeding forms (reviewed in Martín-Durán and Eggers, 2012; Rawlinson, 2014), hatch with a feeding form but settle without food (Kato, 1940; Anderson, 1977) or pass the lobate stage before hatching (Rawlinson et al., 2011; Kato, 1940). It seems worthwhile to investigate whether these cases represent consistent interspecific differences or if some (as reported by Teshirogi et al., 1981 and Bolaños, 2008) might represent poecilogony as described in some other spiralians (e.g. the spionid *Boccardia proboscidea*, Gibson, 1997; or the gastropod *Alderia willowi*, Krug, 2007). An even more interesting possibility would be that the conventional planktotroph hatchling might undergo a change in both morphology and habit without leaving the plankton in response to an environmental factor, specifically food availability. In the case of the Winnie-the-Pooh Müller’s larvae, this possibility seems plausible because a) these larvae transition reversibly from lobate to ovate transiently when disturbed – albeit without altering the arrangement of the ciliary band as they would at metamorphosis – and b) in lab culture we find they undergo irreversible metamorphosis at smaller size when kept unfed (GvD, unpublished observations).

Macrophagous carnivory is clearly not risk-free, especially for a small soft-bodied ciliated blob. While a diet of unicellular algae may require special apparatus and entail risks varying from mere scarcity to outright toxicity or indigestibility, macrophagy requires the predator to encounter and subdue large and possibly well-armored or vigorously-swimming prey. In our observations we noticed several suggestions of potential challenges: for the smallest polyclad larvae, e.g. hatchlings raised in the lab, it seemed difficult for these small larvae to subdue and break through the cuticle of even the smallest prey we offered (of course, it could be that in nature they would choose smaller prey). Furthermore, crustacean appendages in particular are sharp and fast-moving and potentially damage, not just dislodge, these predators; we wonder if this is why we commonly observed polyclads initially attaching to dorsal carapace, where thrashing appendages can’t reach, and riding the swimming prey for some time (e.g. see Fig. 4). Assuming the worms do subdue their prey and break through defenses, engulfing a large amount of tissue in a single feeding – sometimes seemingly more than initial body mass – may impose a significant risk for injury or lead to other vulnerabilities: although we have not measured it, for example, recently-fed worms appeared to swim much more slowly (GvD, pers. obs.). Finally, the prey itself may conceal risks: on several occasions, while feeding lab-reared hatchlings with wild-caught plankton, cultures crashed abruptly, with sick and dying worms joining the carcasses of their prey at the bottom of the bowl (GvD, pers. obs.); we wonder if these events represent toxin bioaccumulation. Despite such risks, it seems likely that macrophagous carnivory constitutes a viable short-cut to planktotrophy: instead of making their own apparatus to collect unicellular food, these flatworms eat someone else who has such devices, using essentially the same feeding apparatus present in adults.

The dichotomy between direct and indirect development has long been associated with numerous ecological and life-history correlates, especially dispersal potential, resource exploitation, and complex trade-offs between parental size, investment, and fecundity (Thorson, 1950; Jägersten 1972; Strathmann, 1985; Marshall et al., 2012). Indirect development, especially if it involves distinct resource use, typically requires developmental specialization for swimming, feeding, growth and even defense ranging from comparatively subtle – e.g., the velum of bivalve veligers, which are otherwise small clams – to extreme, such as the distinct body plan and extreme metamorphosis of planktotrophic echinoderm larvae (Jägersten, 1972; Strathmann, 1985). The pelagic flatworms we studied here, through the strategy of macrophagous carnivory, achieve the ecological status of larvae without the developmental burden of morphological novelty. One might equally say that these species achieve the developmental simplification of direct embryogenesis without the burden of maternal investment. A companion paper provides an equivalent case for hoplonemerteans. These discoveries suggest that macrophagous carnivory might be a more widespread larval strategy than previously recognized.

## Acknowledgements

We thank Svetlana Maslakova for numerous consultations, lab space, and taxonomic advice, Maureen Heaphy for help with sequencing, Gustav Paulay for taxonomic advice and sharing sequence data, and Hiroshi Kajihara for translating passages of Teshirogi et al. (1981). This study was partially funded by São Paulo Research Foundation (FAPESP) grant 2019/10375-8 to CM.

